# A map of binary SARS-CoV-2 protein interactions implicates host immune regulation and ubiquitination

**DOI:** 10.1101/2021.03.15.433877

**Authors:** Dae-Kyum Kim, Benjamin Weller, Chung-Wen Lin, Dayag Sheykhkarimli, Jennifer J. Knapp, Nishka Kishore, Mayra Sauer, Ashyad Rayhan, Veronika Young, Nora Marin-de la Rosa, Oxana Pogoutse, Kerstin Spirohn, Alexandra Strobel, Florent Laval, Patrick Schwehn, Roujia Li, Simin Rothballer, Melina Altmann, Patricia Cassonnet, Guillaume Dugied, Atina G. Cote, Lena Elorduy Vergara, Isaiah Hazelwood, Bingruo B. Liu, Maria Nguyen, Ramakrishnan Pandiarajan, Patricia A. Rodriguez Coloma, Luc Willems, Jean-Claude Twizere, Caroline Demeret, Yves Jacob, Tong Hao, Dave E. Hill, Claudia Falter, Marc Vidal, Michael A. Calderwood, Frederick P. Roth, Pascal Falter-Braun

## Abstract

Key steps in viral propagation, immune suppression, and pathology are mediated by direct, binary, physical interactions between viral and host proteins. To understand the biology of severe acute respiratory syndrome coronavirus 2 (SARS-CoV-2) infection, we generated an unbiased systematic map of binary interactions between viral and host proteins, complementing previous co-complex association maps by conveying more direct mechanistic understanding and potentially enabling targeted disruption of direct interactions. To this end, we deployed two parallel strategies, identifying 205 virus-host and 27 intraviral binary interactions amongst 171 host and 19 viral proteins, and confirming high quality of these interactions via a calibrated orthogonal assay. Host proteins interacting with SARS-CoV-2 proteins are enriched in various cellular processes, including immune signaling and inflammation, protein ubiquitination, and membrane trafficking. Specific subnetworks provide new hypotheses related to viral modulation of host protein homeostasis and T-cell regulation. The binary virus-host protein interactions we identified can now be prioritized as targets for therapeutic intervention. More generally, we provide a resource of systematic maps describing which SARS-CoV-2 and human proteins interact directly.

## INTRODUCTION

Coronavirus disease of 2019 (COVID-19), a severe respiratory disease that emerged in December 2019, is caused by severe acute respiratory syndrome coronavirus 2 (SARS-CoV-2) (Huang et al., 2020; Wu et al., 2020b). As of March 16th, 2021, over 120 million confirmed cases and ~2.7 million deaths have been reported globally (WHO report of coronavirus). Despite intensive research by the scientific community, many important questions regarding molecular viral mechanisms and COVID-19 etiology remain unanswered. Several vaccines against SARS-CoV-2 have been approved, and their deployment has begun in various countries. However, no vaccine offers 100% protection against the disease, vaccination of the global population will take significant time, and multiple SARS-CoV-2 variants have emerged with varying potential to escape vaccine-derived immunity (Grubaugh et al., 2021). Moreover, therapies for infected patients who are acutely ill or suffering from ‘Long COVID’ have had limited efficacy (Carfì et al., 2020). SARS-CoV-2 is likely to be a public health risk into the foreseeable future, thus necessitating a deeper molecular understanding, with the potential to inform effective treatments of infected patients and preventive strategies.

Viruses exploit host processes throughout their replication cycle and must evade or block host immune responses. Several virus-host interactome (‘virhostome’) studies for other viruses have demonstrated the value of systematic network maps for understanding key viral processes (Calderwood et al., 2007; Muller et al., 2012; Neveu et al., 2012; Pfefferle et al., 2011). We have previously shown that systematic virus-host network maps can identify viral pathway perturbations underlying clinical disease manifestations (Gulbahce et al., 2012). For SARS-CoV-2, several studies have identified physical associations between viral and host proteins using affinity purification followed by mass spectrometry (AP-MS) (Gordon et al., 2020a, 2020b; Li et al., 2021; Nabeel-Shah et al., 2020; Stukalov et al., 2020), and identified spatially proximal host proteins using biotin identification (BioID) (Laurent et al., 2020; Samavarchi-Tehrani et al., 2020; St-Germain et al., 2020). Different technologies provide complementary views of the interactome, and protein relationships identified by distinct methods have different biological properties (Yu et al., 2008). Importantly, to this date, no study sought to identify direct interactions between the proteins of SARS-CoV-2 and its human host, as opposed to protein pairs that are indirectly-associated or spatially proximal. Knowledge of direct protein-protein interactions (PPIs) is important for a mechanistic understanding of the virus-host relationship and enables efficient discovery of drugs that can disrupt these interactions (Lu et al., 2020) to interfere with the viral life cycle.

To sensitively identify direct interactions between viral and human host proteins, we carried out protein interaction screens using the yeast two-hybrid (Y2H) system (Fields and Song, 1989), for which the vast majority of identified interactions are known to be direct (or ‘binary’) (Luck et al., 2020; Rolland et al., 2014). In total, we tested >400,000 virus-host protein pairs by each of two binary assay implementations to reveal a network of 205 direct virus-host and 27 intraviral interactions amongst 171 host and 19 viral proteins. Validation of the identified PPIs by an orthogonal split-luciferase based assay, the NanoLuc two-hybrid (N2H) system (Choi et al., 2019), demonstrated high data quality. The resulting binary virhostome interaction map suggested extensive viral targeting of host proteins mediating (i) regulation of immune signaling and inflammation; (ii) ubiquitin-mediated protein regulation and degradation; and (iii) the membrane trafficking. These data comprise an important resource for SARS-CoV-2 research, identifying a largely novel set of direct protein interactions that furthers our mechanistic understanding of SARS-CoV-2 and offers potential entry points for development of targeted COVID-19 therapeutics.

## RESULTS AND DISCUSSION

### Overview of screening methods

Because protein interaction screening methods differ in the subset of interactions they can detect (Braun et al., 2009), we employed two independent Y2H versions (**Fig. 1A**): (i) an auxotrophic imidazoleglycerol-phosphate dehydratase (“HIS3”) selection with a growth-based readout (Y2H_HIS3_) enabled by interaction-dependent transcriptional activation of the *HIS3* reporter gene, a system which has been used previously to generate high-quality interactome network maps (Altmann et al., 2018, 2020; Rual et al., 2005; Yu et al., 2008); and (ii) a green fluorescent protein (GFP) system (Y2H_GFP_) based on the Barcode Fusion Genetics (BFG)-Y2H technology (Yachie et al., 2016), in which interaction-dependent activation of the GFP reporter gene is measured by fluorescence-activated cell sorting (FACS), enabling sorting of GFP-positive cells from a pooled culture (Kim et al., 2021; Yachie et al., 2016). The complementary detection profiles of Y2H_HIS3_ and Y2H_GFP_ (see below) are achieved by different system configurations: Y2H_HIS3_ is based on N-terminally tagged viral and human open reading frame (ORF) constructs expressed from low-copy plasmids, while Y2H_GFP_ employs C-terminal activation domain (AD) fusions of human ORFs expressed from high-copy plasmids, and also differs in terms of the length and sequence of the linker between the ORF and AD sequences.

**Figure 1.**
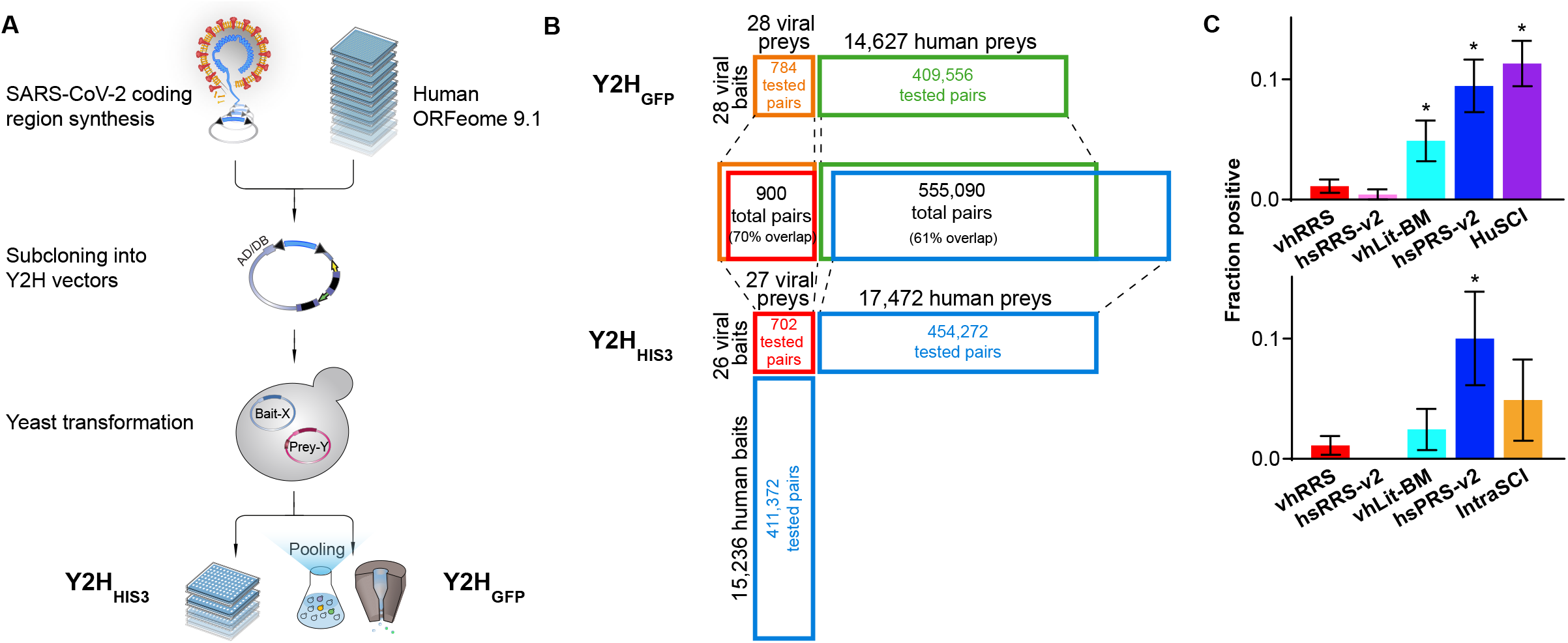
Generation and quality assessment of proteome-wide direct interaction maps amongst SARS-CoV-2 and human host proteins. **(A)** Schematic description of the experimental pipeline. **(B)** Screening space for each of the two parallel Y2H_HIS_ and Y2H_GFP_ screens. “Percent overlap” is relative to the union of protein pairs tested by both methods. **(C)** Orthogonal N2H assay validation of HuSCI and IntraSCI along with positive (hsPRS-v2 and vhLit-BM) and negative (hsRRS-v2 and vhRRS) benchmarking sets. Asterisks indicate significant differences from viral negative benchmarking set (*p* < 0.05, Fisher’s exact test).

### Clone assembly and construction

Clones encoding SARS-CoV-2 proteins were independently synthesized and assembled (**top panel of Fig. 1A**). For Y2H_HIS3_, Gateway Entry clones were generated based on SARS-CoV-2 reference (non-codon-optimized) ORFs from the sequenced genomes of three viral isolates (Wu et al., 2020a) (**see Methods**). Viral ORFs were transferred by recombinational cloning to the appropriate Gateway Destination vectors, which respectively express the Gal4 DNA binding domain (DB) fused to the N-terminus of the ‘bait’ protein (DB-X), and the Gal4 AD fused to the N-terminus of the ‘prey’ protein (AD-Y) (**see Methods**). Human ORFs from two previously reported collections of haploid yeast clones were used, expressing ORFs N-terminally fused to Gal4 DB and AD domains (Luck et al., 2020).

For Y2H_GFP_, our codon-optimized SARS-CoV-2 ORF collection was cloned into Gateway Entry plasmids (Kim et al., 2020), and then transferred into barcoded low-copy bait and prey plasmids as N-terminal fusions. To interrogate human ORFs, a collection of high-copy prey plasmids with C-terminal fusions (**see Methods**) was assembled that collectively covered ~14K human ORFs, each generally being represented in two distinct uniquely-barcoded plasmids. We previously reported a subset of ~14K of these plasmids, with one uniquely-barcoded plasmid per ORF (Luck et al., 2020), and an additional set of plasmids for ~14K ORFs was generated for this study to provide independently-barcoded replication.

For each of the two screens, the independently-cloned viral ORFs included: non-structural proteins (NSPs) 1-16, derived from ORF1AB; the four structural proteins spike (S), envelope (E), membrane (M) and nucleocapsid (N); and 9 accessory genes, ORF3A, 3B, 6, 7A, 7B, 8, 9B, 9C and 10 (**Supplementary Table 1A; see Methods**). Because the genome sequence and annotation of viral ORFs used for the two cloning efforts (Chan et al., 2020; Wu et al., 2020a) diverged for NSP12, ORF3B and ORF10, the two clone sets together cover 30 viral ORFs. Therefore, we hereafter use a recently proposed nomenclature (Jungreis et al., 2020), so that NSP12 (Chan et al., 2020) and NSP12 (Wu et al., 2020a) are referred to as NSP12-1 and NSP12-2, respectively, and ORF3B (Chan et al., 2020) and ORF3B (Wu et al., 2020a) are referred to as ORF3D and ORF3B, respectively (**Supplementary Table 1A**).

Screening for binary interactions between viral and human host proteins and amongst viral proteins Using Y2H_HIS3_, 27 viral ORFs were screened against 17,472 human ORFs (83% ‘completeness’, i.e., covering 83% of the potential search space) in both orientations so that each protein is screened as both bait and prey (**Fig. 1B**). Screens proceeded by mating each bait strain to ‘mini-pools’ of prey strains, picking specifically growing colonies, subjecting these to secondary phenotyping, and sequencing of plasmids from positive colonies to identify interacting proteins (**see Methods**). Each identified human interaction candidate was tested three times in pairwise Y2H experiments against all 27 viral ORFs to verify the candidates. Only pairs scoring positive at least two times and not exhibiting an ‘auto-activation’ phenotype (growth in selective media in the absence of the prey plasmid) were considered *bona fide* Y2H interactions (**see Methods**). This screen yielded 119 interactions, collectively involving 14 viral and 93 human host proteins. We refer to this binary human SARS-CoV-2 interactome map as HuSCI_HIS3_ (Human-SARS-CoV-2 interactome via the Y2H_HIS3_ system).

For the Y2H_GFP_ screen, we examined 14,627 prey human ORFs (70% completeness), most of which were represented by two uniquely-barcoded plasmids, collectively corresponding to 27,671 uniquely-barcoded yeast strains. Each of these strains was screened against bait strains for 28 viral ORFs (each represented by 2-6 uniquely-barcoded strains, collectively corresponding to 82 uniquely barcoded bait strains). Thus, we screened 409,556 bait-prey combinations represented by 2,269,022 uniquely barcoded diploid strains (**Fig. 1B**). Barcode sequencing of GFP-positive cells allowed quantitative assessment of auto-activity levels and therefore, unlike Y2H_HIS3_, enabled identification of genuine PPIs even for high-background baits via barcode enrichment analyses. After stringent filtering (based on effect size, significance, and replicate agreement) and retesting using the *HIS3* marker, the Y2HGFP screen identified 93 interactions among 13 viral and 84 human host proteins. We refer to this binary human SARS-CoV-2 interactome map as HuSCIGFP (Human-SARS-CoV-2 interactome via the Y2H_GFP_ system), and refer to the union of HuSCI_HIS3_ and HuSCI_GFP_ as HuSCI (**Supplementary Table 1B**). The Y2H_GFP_ screen also yielded 27 intraviral interactions amongst 19 viral proteins (**Supplementary Table 1C**), here termed as IntraSCI (intraviral SARS-CoV-2 interactome).

Having collectively identified 205 direct virus-host and 27 intraviral interactions amongst 171 host and 19 viral proteins, we proceeded to assess the quality of these candidate interactions.

### Assessment of interaction data quality

As a first level of quality control, we examined the overlap of interactions among our datasets. Considering that ~60% of all viral / human protein pairs were assayed in both screens, and that each assay has an assay sensitivity (fraction of true interactions that an assay can detect in a fully-saturated screen) of 20-25%, and an estimated sampling sensitivity (the extent to which each screen is saturated) of 50% and 60% for the two screens, we could have expected ~3% overlap between the screens *a priori,* which is close to the observed 3.6% overlap (**see Methods**). Given these screening parameters, we estimate that our merged dataset covers about 20% of all binary SARS-CoV-2 virus host interactions (**see Methods**), which is comparable to previous high quality binary interactome datasets (Luck et al., 2020; Rolland et al., 2014; Yu et al., 2008).

To further assess data quality experimentally, we deployed an established empirical framework based on validation with a well-calibrated orthogonal biochemical interaction assay (Choi et al., 2019; Luck et al., 2020; Rolland et al., 2014; Yu et al., 2008). Because any interaction assay can detect only a subset of *bona fide* interactions (Braun et al., 2009), the fraction of pairs validated in an orthogonal assay (the ‘validation rate’) must be calibrated against a positive control set of well-documented interactions (Positive Reference Set, PRS) and a negative control set of randomly-selected protein pairs (Random Reference Set, RRS). We used the previously established human PRS / RRS version 2 (hsPRS-v2 and hsRRS-v2) sets as human positive and random reference sets for calibration (Braun et al., 2009; Choi et al., 2019). As another benchmark, we derived a collection of 55 human coronavirus / host protein interactions from the literature (Cusick et al., 2009) using the criteria of being supported by multiple sources, of which at least one indicates a binary interaction. We refer to this benchmark as the virus-host literature binary multiple reference set (vhLit-BM; **Supplementary Table 2A**). We further established a virus-host Random Reference Set (vhRRS) of SARS-CoV-1 and 2 viral and human host protein pairs by randomly selecting 180 protein pairs not previously reported as interactions.

We subjected HuSCI, IntraSCI, and each benchmark set of protein pairs to the orthogonal yeast-based N2H validation assay (yN2H; **Fig. 1C and Supplementary Fig. 1**) (Choi et al., 2019). We set stringent thresholds for yN2H such that only one vhRRS and no hsRRS-v2 pair scored positive. At this threshold, HuSCI alone and HuSCI merged with IntraSCI exhibited validation rates of 11 ± 2% and 10 ± 2%, respectively, which were statistically indistinguishable from the two positive control sets (hsPRS-v2 and vhLit-BM, 9 ± 2%; *p* = 0.88 and 5 ± 2%, *p* = 0.06; Fisher’s exact tests relative to the union of HuSCI and IntraSCI), and significantly better than negative control set validation rates (hsRRS-v2, 0.4 ± 0.4%, *p* = 4 × 10^−7^ and vhRRS 1.1 ± 0.6%; *p* = 1 × 10^−7^; Fisher’s exact tests relative to the union of HuSCI and IntraSCI). IntraSCI contained too few interactions for a separate statistical evaluation. Thus, our virus-host interaction map (HuSCI, **Supplementary Table 1B** together with IntraSCI, **Supplementary Table 1C**) exhibited biophysical quality at least *on par* with human-human and virus-host benchmark sets of interactions that have been supported by multiple experiments in the manually curated literature.

Thus, we present a validated high-quality network of 205 virus-host and 27 intraviral direct, binary interactions amongst 171 host and 19 viral proteins (**Fig. 2A**).

**Figure 2.**
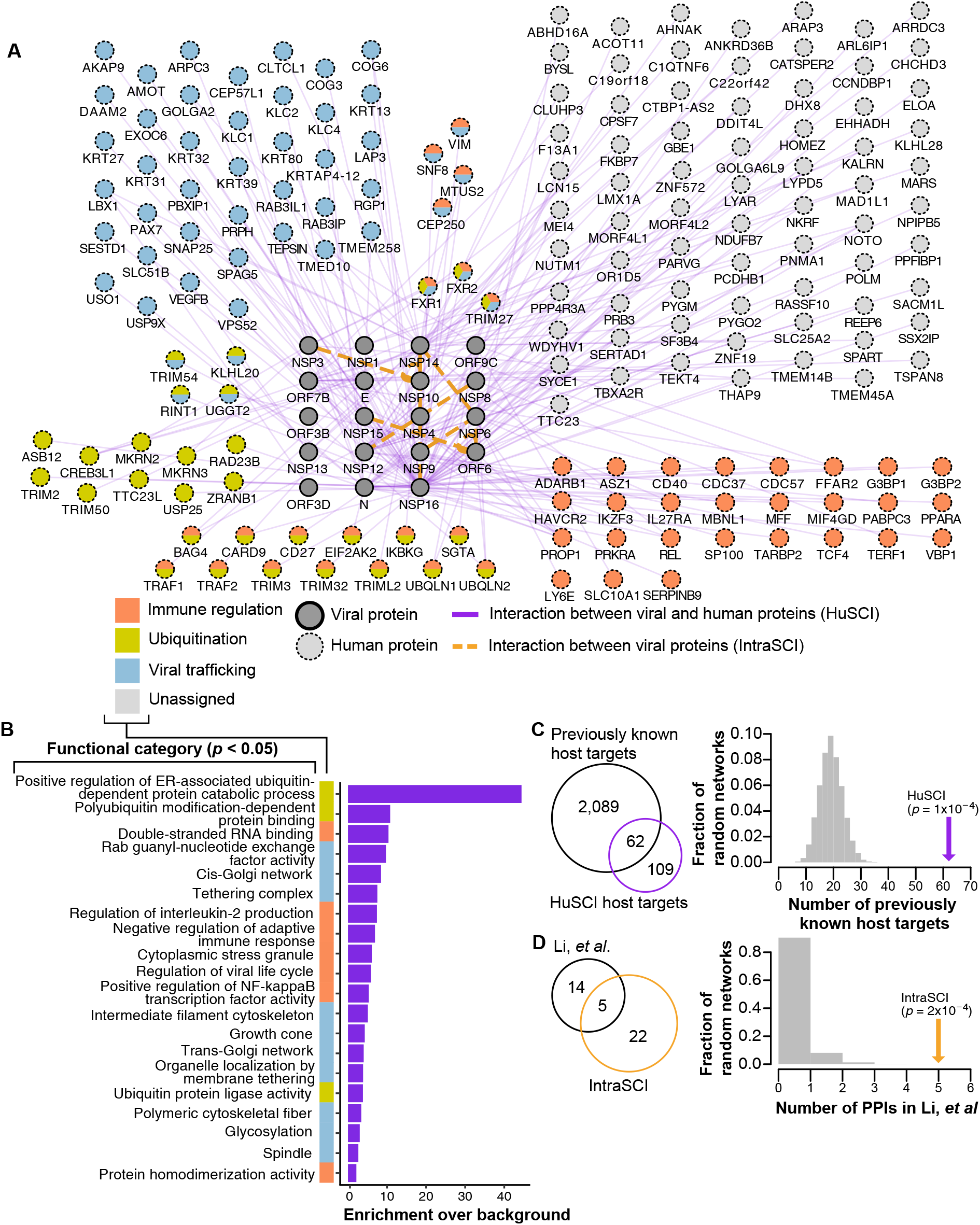
Network representation and functional assessment of proteome-wide direct interaction map between SARS-CoV-2 and human host proteins. **(A)** The integrated HuSCI and IntraSCI networks. Host proteins grouped by the most enriched broad functional terms, including immune regulation, ubiquitination and viral trafficking. **(B)** Functions enriched amongst the host proteins found in HuSCI. **(C)** Comparison of previously identified targets of other viruses with SARS-CoV-2 targets identified in HuSCI. **(D)** Comparison of previously known intraviral interactions and IntraSCI.

The high quality of our dataset was further supported by the finding that host proteins detected as viral interaction partners in our screen were enriched with Gene Ontology terms related to viral infection (**Fig. 2B**), and by the observation that 62 of the here-identified viral interactors are known targets of other viruses (Orchard et al., 2014) (*p* = 1 × 10^−4^ by empirical testing) (**Fig. 2C**). Moreover, we find that a significant number of physically targeted proteins in HuSCI have been found to change phosphorylation status upon SARS-CoV-2 infection (*p* = 0.002 by empirical testing; **Supplementary Fig. 3**) (Bouhaddou et al., 2020; Stukalov et al., 2020). In addition, our IntraSCI network of 27 interactions showed a significant overlap of 5 interactions with a previous set of 19 intraviral interactions (Li et al., 2021) (*p* = 2 × 10^−4^ by empirical resampling) (**Fig. 2D**). These observations together suggested that our map is of high biophysical quality, and enriched for host proteins that are relevant to the biology of SARS-CoV-2.

### Complementarity of Y2H and AP-MS datasets

All previous maps of association between SARS-CoV-2 and host proteins (Gordon et al., 2020a, 2020b; Li et al., 2021; Nabeel-Shah et al., 2020; Stukalov et al., 2020) have been generated using methods that cannot differentiate between direct and indirect interactions. By contrast, binary assays such as Y2H detect protein interactions that are predominantly direct (Luck et al., 2020; Rolland et al., 2014). This fundamental difference necessarily reduces the expected overlap between protein pairs identified in these assays (Yu et al., 2008). More surprising is the overall low agreement (~2%) observed between different pairings of these four AP-MS based association datasets, although this may be partially explained by differences in experimental setup, i.e. distinct cell lines and protocols (Braun, 2013).

Non-overlap between different association maps (and between direct interaction and association maps) might also be expected if the SARS-CoV-2 virus-host interactome contains many weak interactions that are less reliably detected. Interestingly, the validation rates of known virus-host interactions (vhLit-BM) and the random reference set of virus-host pairs (vhRRS) are less clearly separated than the corresponding positive and random human-human protein interaction benchmarks (**Supplementary Fig. 1**). Although it is possible that this phenomenon is due to imperfect quality of our literature-derived benchmark for coronaviral-host interactions, the yN2H data points to a higher prevalence of interactions amongst randomly chosen virus-host pairs (vhRRS), which may in turn stem from limited evolutionary selection against non-specific interactions. Previous systematic studies of interactions between yeast and human proteins observed an overall density of between-species interactions that was comparable to that of within-species interactions, despite the billion-year absence of selection to maintain yeast-human interactions (Zhong et al., 2016). This suggested that, in the evolution of protein interactions, selection against non-specific protein interactions may be as important as selection for specific protein interactions. A similar conclusion was reached by an earlier study on a single yeast SH3 binding peptide, which interacted very specifically with a single domain in its native proteome, but showed a broad interaction profile with SH3 domains from other organisms (Zarrinpar et al., 2003). Thus, the fact that SARS-CoV-2 has only recently jumped a species barrier is consistent with an increased abundance of less-reliably detected (presumably weaker) virus-host protein interactions.

Despite fundamental assay differences, low overlap between AP-MS datasets, and a potential tendency for SARS-CoV-2 to have weaker interactions with human host proteins, there were many overlaps between our network and protein pairs reported in other studies. Among our 205 HuSCI virus-host interactions, 11 (~5.4%) had been previously reported by at least one SARS-CoV-2 AP-MS study (Gordon et al., 2020a, 2020b; Li et al., 2021; Nabeel-Shah et al., 2020; Stukalov et al., 2020) (**Supplementary Table 1B**). Additionally, we found 29 human proteins in common with four previous AP-MS studies (Gordon et al., 2020a, 2020b; Li et al., 2021; Nabeel-Shah et al., 2020; Stukalov et al., 2020), albeit targeted by other viral proteins than the previously-reported (potentially indirect) associations. The interactions most consistently identified in other datasets are between the viral N protein and host stress granule proteins G3BP1 and G3BP2. Previously, it was shown that overexpression of the viral N protein stimulates stress granule formation (Gupta et al., 2017; Reineke and Lloyd, 2015). The fact that all interactome datasets identify the interaction of N with G3BP1 and G3BP2 (Nabeel-Shah et al., 2020; Samavarchi-Tehrani et al., 2020), increase the confidence that this pathway is targeted. While differences were expected between the specific interactions and associations of different studies, we do find overall agreement at the functional level (**see next section**), in keeping with previous reports that Y2H and AP-MS reveal complementary maps but mutually consistent perspectives on the interactome.

### Functions enriched in a SARS-CoV-2/human virhostome network

To identify which host functions are targeted by SARS-CoV-2, we performed GO enrichment analysis (**see Methods**). The terms most enriched amongst host proteins can be categorized more broadly into three functional groups: (i) immune regulation, (ii) ubiquitination and (iii) viral trafficking (**Fig. 2A and B**). Comparison with the four AP-MS based networks revealed substantial functional overlap, especially for terms related to viral trafficking, including vesicle-mediated transport to the plasma membrane and Golgi network (**Supplementary Fig. 2**). Despite functional consistency between the different datasets, the divergence between the interactions we identified and the previously reported associations illustrates the complementarity of these methods and emphasizes the advantage of using both approaches to understand SARS-CoV-2 biology. Moreover, in addition to confirming previously identified cellular processes, our network revealed novel functional enrichment that may shed light on viral modulation of immune regulation and virally-targeted ubiquitination processes.

The most obviously clinically-relevant functions of viral targets relate to immune regulation. Infection with SARS-CoV-2 is often accompanied by dysregulation of interferon (IFN)-mediated antiviral response, as well as elevated inflammatory cytokine signaling (Blanco-Melo et al., 2020; Hadjadj et al., 2020; Karki et al., 2021). Dysregulation of IFN responses facilitates viral replication in the early stages of infection, while the so-called cytokine storm leads to acute respiratory distress syndrome in severe cases of COVID-19 (Fajgenbaum and June, 2020). Recent studies suggest that SARS-CoV-2 inhibits type I IFN induction by blocking host mRNA translation (Thoms et al., 2020), possibly by targeting TBK1 (Gordon et al., 2020a, 2020b). Additionally, the transcriptional activation of IFN-stimulated genes is inhibited by preventing STAT1 nuclear import by an ORF6-NUP98 interaction (Miorin et al., 2020).

Members of the tripartite motif (TRIM) protein family of E3 ligases, which act to ubiquitinate target proteins, have been implicated in regulating antiviral host defenses and innate immune signaling (van Gent et al., 2018). We found several members of the TRIM protein family, namely TRIM2, 3, 27, 32, 50 and 54, to be interaction targets of viral NSP16 and NSP14. TRIM27 has been shown to modify TBK1 and IKBKG, thereby modulating activation of IRF3/IRF7 (Zheng et al., 2015) and NF-κB (Zha et al., 2006), which, in turn, activate transcription of type I IFN and proinflammatory cytokines, respectively. Interestingly, TRIM27 was also found to be a degradation target of ICP0 during HSV-1 infection (Conwell et al., 2015). TRIM32 modulates innate immune responses in several ways. It negatively regulates TRIF and thereby TRL3/4 responses (Yang et al., 2017), as well as type I IFN production through ubiquitination of STING (Zhang et al., 2012). TRIM32 was further shown to inhibit influenza A replication by targeting its polymerase for proteasomal degradation (Fu et al., 2015). The direct NSP16-TRIM32 interaction we found was previously reported as an association (Gordon et al., 2020b). Additionally, we find an interaction between NSP14 and IKBKG, as well as multiple interactions between the NF-kB family member REL and NSP14, NSP16, and NSP9. Taken together, the identified interactions reveal multiple direct routes by which SARS-CoV-2 may be targeting both the type I IFN pathway, and hence antiviral host innate immune signaling, and inflammatory cytokine signaling.

A phenomenon observed in COVID-19 patients, for which the mechanism is unclear, is the functional exhaustion of cytotoxic lymphocytes during the adaptive immune response (Zheng et al., 2020). Here, we find that host proteins related to immune response are enriched amongst the partners of viral NSP6 (*p* < 10^−2^ by empirical testing), as are membrane proteins (*p* < 10^−2^ by empirical testing). Supporting the latter enrichment, NSP6 is itself a membrane protein that induces formation of viral double-membrane vesicles (Angelini et al., 2013). We find that NSP6 interacts with membrane regulators of cytotoxic lymphocytes responding to virus infection, including CD40 (Bennett et al., 1998), CD27 (Ochsenbein et al., 2004) and IL27RA (Wehrens et al., 2018). CD40’s interaction with its ligand CD40L is important for T cell priming (Toes et al., 1998) and CD40 deficiency is known to inhibit T cell development against influenza virus (Lee et al., 2003). Use of CD27 agonists is exploited in cancer immunotherapy to co-stimulate the T-cell response (van de Ven and Borst, 2015) and CD27 expression is up-regulated in HIV-infected patients (Ochsenbein et al., 2004). IL27, the ligand of IL27RA, suppresses T cell cytotoxicity and viral control during cytomegalovirus infection (Wehrens et al., 2016). Thus, our map points to NSP6 as a potential regulator of T cell development and related COVID-19 symptoms. More broadly, our map is enriched for host proteins relevant to immune regulation and provides numerous mechanistic hypotheses.

The second functional category we identified is related to ubiquitination and suggests functional cross-talk between different targeted pathways. In addition to the ubiquitination-dependent modulation of host immune responses discussed above, we find ubiquitin-dependent degradation to be targeted by SARS-CoV-2. Hijacking the ubiquitin proteasome system is a common trait of almost all viruses (Banks et al., 2003; Fanunza et al., 2019; Tang et al., 2018). Supporting the idea that this is occurring for SARS-CoV-2, we found several viral proteins to target host proteins involved in ubiquitin-mediated degradation. More specifically, we identified viral ORF3D, ORF6 and ORF9C as interacting with the host proteins UBQLN1/2. This is particularly interesting in light of the recent discovery that ORF9C attenuates antiviral response in lung epithelial cells in a proteasome-dependent manner (Andres et al., 2020). Human UBQLN1/2 mediates protein degradation of its interaction partners, e.g. hnRNPA1 (Gilpin et al., 2015) and TDP43 (Cassel and Reitz, 2013), by recruiting them to the 26S protease or autophagosome (Renaud et al., 2019).

A third functional category that was strongly represented among our interactors is related to viral trafficking via the ER-Golgi membrane network. Several similar terms were described by the physical association studies including: endomembrane system organization (Gordon et al., 2020a, 2020b), Golgi membrane (Stukalov et al., 2020), and Golgi vesicle transport (Li et al., 2021). The robust identification of these processes by all experimental approaches emphasize their importance in the viral life cycle. Of particular interest was NSP16, which is known to methylate mRNA to facilitate viral replication and escape from innate immune recognition (Wang et al., 2015), but also interacts with many proteins related to viral transport. In addition, we found NSP16 interactors RAB3IL1 (Wandinger-Ness and Zerial, 2014), VPS52 (Conibear et al., 2003), COG6 (Blackburn et al., 2019), and EXO6 (Boehm et al., 2017), which are important factors of transport into the extracellular space via ER-Golgi network, and are closely related to viral trafficking (Sicari et al., 2020). Therefore, in addition to its functions in viral replication, analysis of interactions suggests a role for NSP16 in virion production and release.

### Pairwise test of published SARS-CoV-1 interactions supports conserved immune regulation mechanism

Because interactions are often conserved between species (the ‘interolog’ phenomenon) (Matthews et al., 2001; Walhout et al., 2000; Yu et al., 2004), we sought to cast a wider net by carrying out pairwise assessment of SARS-CoV-2 viral-human pairs corresponding to interactions previously reported for SARS-CoV-1. Of the known SARS-CoV-1 binary virus-host PPIs that had been reported by Y2H, 10% (6 out of 62 PPIs, excluding 14 which are not amenable to testing given due to viral bait autoactivation) were recapitulated in a pairwise HIS3-based Y2H assay (**see Methods**) when substituting SARS-CoV-2 proteins for the originally reported SARS-CoV-1 proteins. The identified interologs included NSP3-MKRN3, NSP8-H2AFY2, NSP8-TERF1, NSP9-FAHD1, NSP13-N4BP2L2, and ORF7B-CAMLG. Two of these interactions, NSP8-TERF1 and ORF7B-CAMLG, were also detected in our HuSCI network. Furthermore, most host proteins in the interologs are associated with functional pathways that were found to be enriched in our network analysis (**see Methods**). The proteins MKRN3 (Kanber et al., 2009) and CAMLG (Peng et al., 2010) are involved in protein ubiquitination. The proteins TERF1 (de Lange, 2005), H2AFY2 (Zhang et al., 2005) and N4BP2L2 (Salipante et al., 2009) are involved in various pathways regulating transcriptional mechanisms. Of particular note is the interaction between viral NSP13, a highly conserved helicase (Chan et al., 2020) that has been associated with suppression of IFN production and signalling (Yuen et al., 2020), and human N4BP2L2. Human N4BP2L2 is known to interact with both neutrophil elastase and the transcriptional repressor GFI1 to modulate the production of neutrophils, a type of white blood cells essential to host innate immunity (Salipante et al., 2009). Interestingly, NSP13 has been suggested to act as a transcriptional regulator, adding to its importance in viral function (Gordon et al., 2020a, 2020b). Taken together, a closer investigation of the direct interaction between viral NSP13 and host N4BP2L2 proteins may be of importance to understanding the mechanisms behind an ‘under siege’ host immune response following SARS-CoV-2 viral infection.

### Limitations and further directions

All protein interaction assays have limitations intrinsic to each method. Y2H assays are limited by the fact that proteins are exogenously expressed with functional assay tags and targeted to the nucleus. The heterologous nature of the assays and circumvention of physiological transcriptional regulation are limitations, but also a benefit in that they enable detection of interactions that might otherwise be missed. For example, screens that rely on the expression of one or both partners in a given cell line and growth condition might miss interactions for proteins that are not expressed in that cell line, even where these interactions are important in other tissues (Hikmet et al., 2020). Despite these limitations, it has been demonstrated repeatedly that Y2H systems, when quality controlled by orthogonal validation with empirically benchmarked assays as done here, yield high-quality interactions that enable important and robust biological insights (Altmann et al., 2020; Choi et al., 2019; Luck et al., 2020; Rolland et al., 2014; Yu et al., 2008).

A known issue with every carefully conducted interaction assay is that true interactions can be missed, with only 20-40% of reference interactions being detectable by any single assay (Braun et al., 2009; Choi et al., 2019; Luck et al., 2020; Yu et al., 2008). This was a major motivation for applying complementary parallel approaches in this study. Future efforts might expand the barcoded ORFeome approach to distinct assay versions, including alternative linkers and orientations for fused bait and prey tags. In addition, assay sensitivity can be increased by implementing the same assay version in different conditions (Kim et al., 2021; Liu et al., 2020). More physiologically relevant protein interactions might also be discovered by conducting our screen in the presence of a third viral protein which might be necessary for mediating other direct interactions. Our genome-wide viral and human barcoded ORFeomes are readily transferable to new genetic and environmental backgrounds.

One of the main advantages of Y2H, unlike protein associations identified by AP-MS, is that the interactions it reports are nearly always direct (Luck et al., 2020; Rolland et al., 2014). None of the viral-host protein association maps generated previously can distinguish between direct and indirect interactions. Knowledge of direct interactions is necessary for an accurate mechanistic understanding of viral infection and progression for potential specific and efficient interventions. A now-possible extension of the Y2H_GFP_ method could exploit its ability to measure many protein pairs in a single pool for rapid screening of potential interaction-disrupting drugs against hundreds of virus-host protein interactions, such as those identified here. Taken together, we expect that combining our map of direct SARS-CoV-2 protein interactions with previously established protein physical association and proximity networks will broaden our mechanistic understanding of viral proliferation and enable rapid development of therapeutic approaches to combat current and future pandemics.

---

**Supplementary Figure 1.**
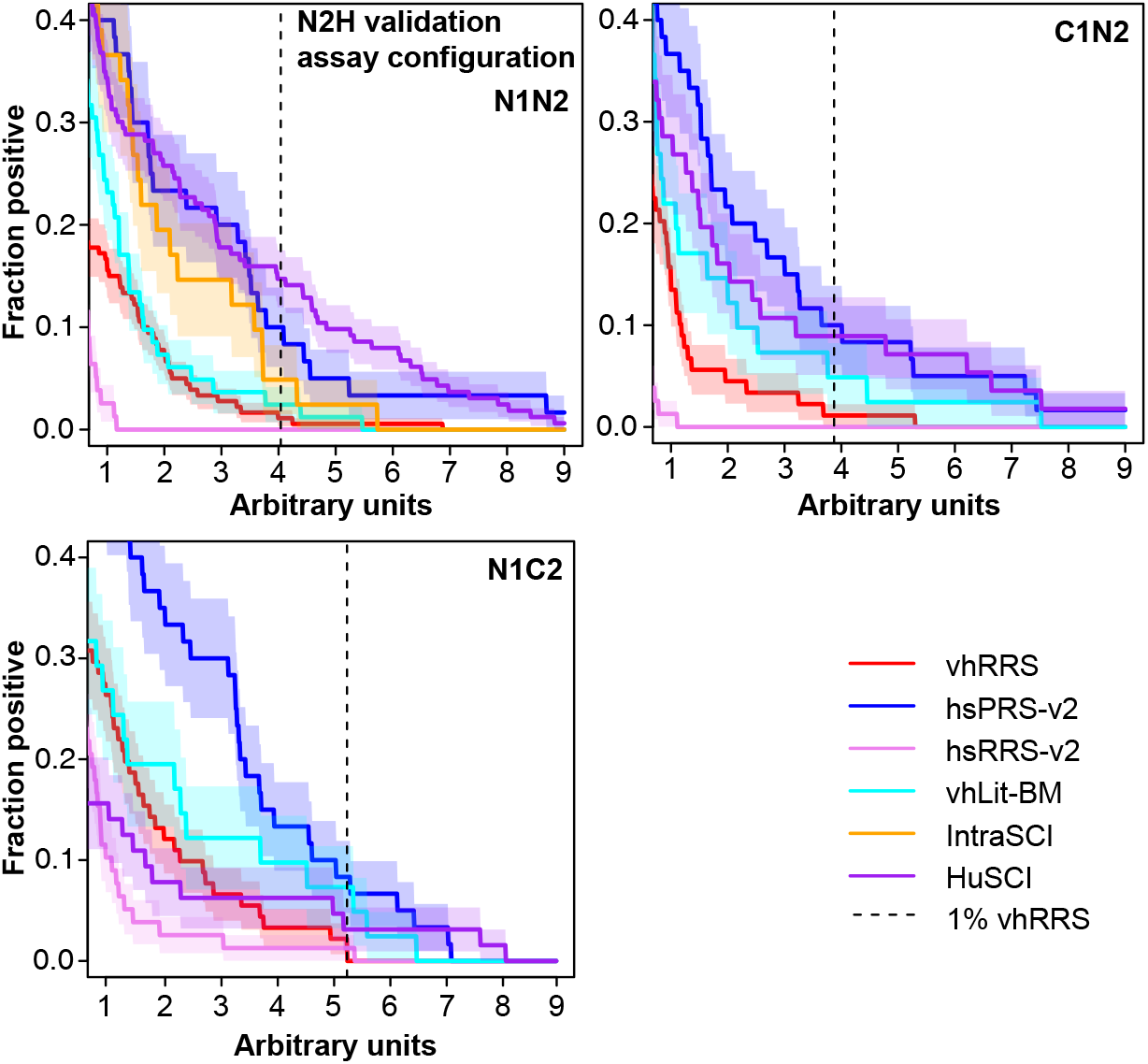
Rate of detection in an orthogonal N2H assay at different scoring thresholds. Rate at which interactions are detected by yN2H for HuSCI and IntraSCI, as well as positive (hsPRS-v2 and vhLit-BM) and negative (hsRRS-v2 and vhRRS) benchmark sets, across stringency thresholds.

**Supplementary Figure 2.**
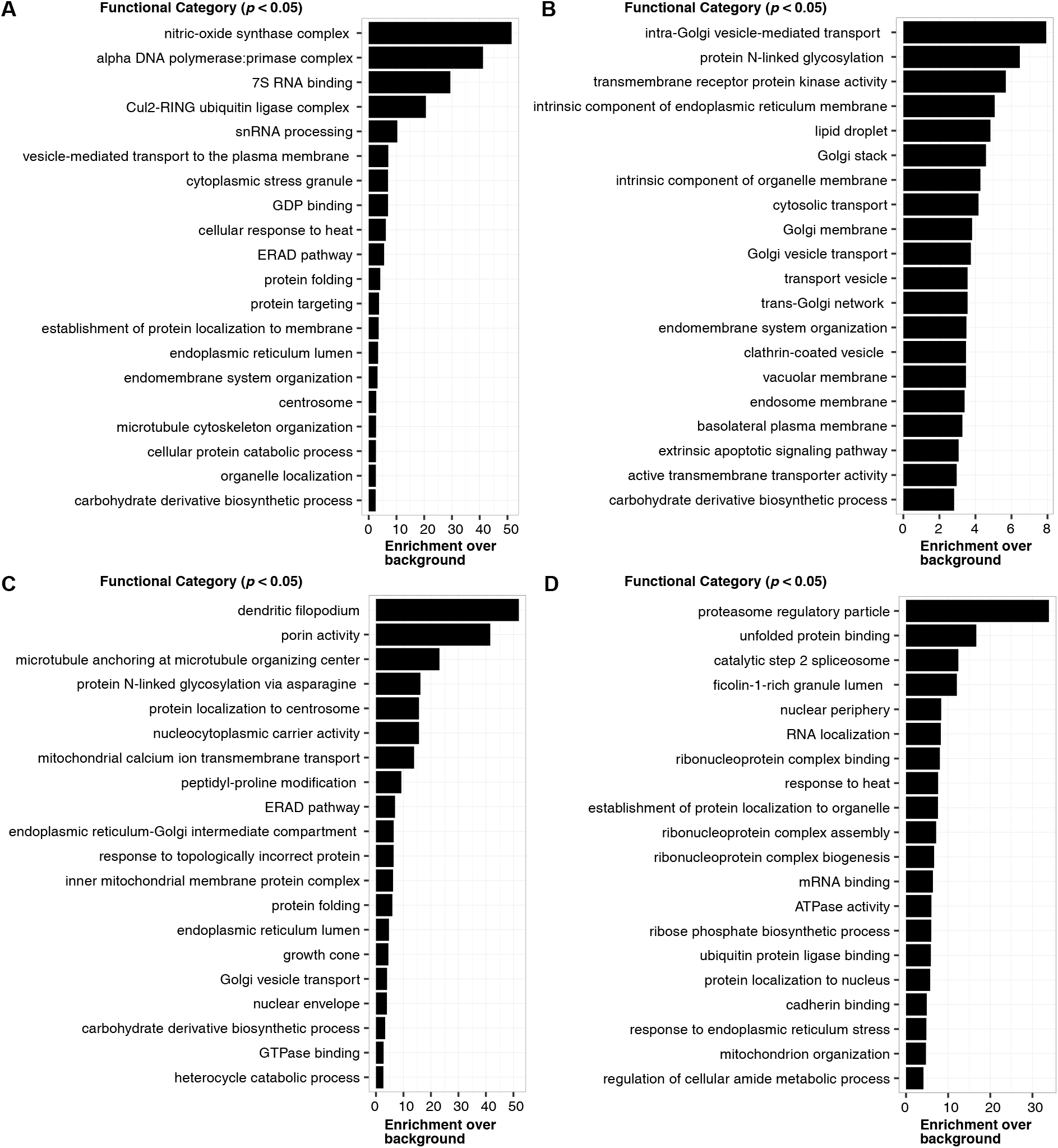
Functional enrichment in the host proteins of four AP-MS based networks. **(A)** Gordon et al. (Gordon et al., 2020a, 2020b) **(B)** Stukalov et al. (Stukalov et al., 2020) **(C)** Li et al. (Li et al., 2021) **(D)** Nabeel et al. (Nabeel-Shah et al., 2020)

**Supplementary Figure 3.**
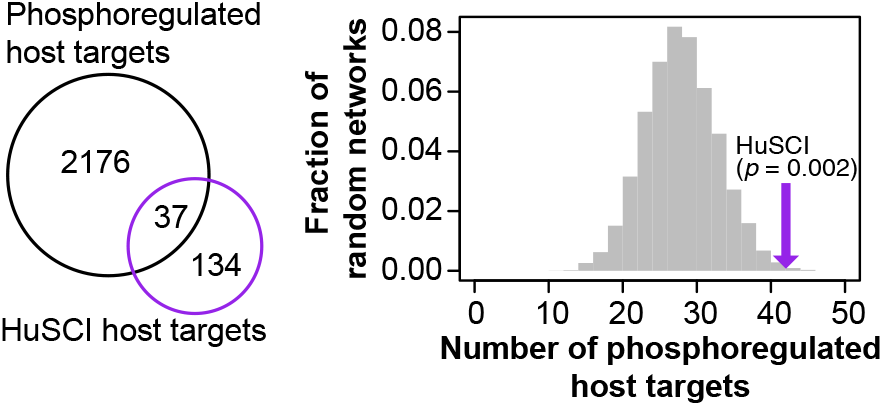
Host targets identified in HuSCI are enriched in differentially phosphorylated proteins following infection by SARS-CoV-2.

## STAR METHODS

- **KEY RESOURCES TABLE**
- **LEAD CONTACT AND MATERIALS AVAILABILITY**
- **EXPERIMENTAL MODEL AND SUBJECT DETAILS**
- **METHOD DETAILS**

○ Selection & clone preparation of functional coding regions from SARS-CoV-2
○ Screening based on HIS3 to generate the HuSCI_HIS_3 map
○ Screening based on GFP to generate the HuSCI_GFP_ map
○ Validation by an orthogonal yN2H assay
○ Bioinformatic analysis of binary interactome map
○ Identification of interologs via pairwise Y2H testing
- **DATA AND CODE AVAILABILITY**

○ The read counting based on the expected barcode pipeline is available at https://github.com/RyogaLi/BFG_Y2H/.
- **QUANTIFICATION AND STATISTICAL ANALYSIS**

## TABLE LEGENDS

**Supplementary Table 1A.** Total list of viral proteins used in both Y2H screens. Annotations are given for the putative function of individual proteins, as well as amino acid sequences.

**Supplementary Table 1B.** Total list of interactions between viral and host proteins in HuSCI. The screen in which the interaction was found (HuSCI_HIS3_ and/or HuSCI_GFP_) is specified, as well as interactions also found by the four AP-MS studies (Gordon et al., 2020a, 2020b; Li et al., 2021; Nabeel-Shah et al., 2020; Stukalov et al., 2020)

**Supplementary Table 1C.** Total list of PPIs amongst viral proteins (IntraSCI) identified by Y2H_GFP_. The overlap with a previous intra-viral PPI study is also indicated (Li et al., 2021)

**Supplementary Table 2A.** List of PPIs in virus-host literature binary multiple reference set (vhLit-BM). The number of methods by which the interaction was identified is indicated.

**Supplementary Table 2B.** List of protein pairs in viral human Random Reference Set (vhRRS).

**Supplementary Table 3A.** Total list of viral clones in AD-Nterm-Cen Y2H destination vector for Y2H_GFP_.

**Supplementary Table 3B.** Total list of viral clones in DB-Nterm-Cen Y2H destination vector for Y2H_GFP_.

**Supplementary Table 4.** Genotypes of toolkit strains used in Y2H_GFP_.

**Supplementary Table 5.** A curated list of previously identified binary interactions between SARS-CoV-1 and human host proteins.

## ACKNOWLEDGEMENTS

This work was supported by a Canadian Institutes for Health Research Foundation Grant (F.P.R.); the Canada Excellence Research Chairs Program (F.P.R.); the Thistledown Foundation (F.P.R.); the LabEx Integrative Biology of Emerging Infectious Diseases (10-LABX-0062; Y.J.) and Platform for European Preparedness Against (Re-)emerging Epidemics, EU (602525; Y.J.); the European Union’s Horizon 2020 Research and Innovation Programme (Project ID 101003633, RiPCoN; P.FB, C.B., P.A.). F.L. was supported by a Belgian American Educational Foundation Doctoral Research Fellowship and a Wallonia-Brussels International (WBI)-World Excellence Fellowship. M.V. is a Chercheur Qualifié Honoraire from the Fonds de la Recherche Scientifique (FRS-FNRS, Wallonia-Brussels Federation, Belgium).

## REFERENCES

Altmann, M., Altmann, S., Falter, C., and Falter-Braun, P. (2018). High-quality yeast-2-hybrid interaction network mapping. Curr. Protoc. Plant Biol. 3, e20067.

Altmann, M., Altmann, S., Rodriguez, P.A., Weller, B., Elorduy Vergara, L., Palme, J., Marín-de la Rosa, N., Sauer, M., Wenig, M., Villaécija-Aguilar, J.A., et al. (2020). Extensive signal integration by the phytohormone protein network. Nature 583, 271–276.

Andres, A.D., Feng, Y., Campos, A.R., Yin, J., Yang, C.-C., James, B., Murad, R., Kim, H., Deshpande, A.J., Gordon, D.E., et al. (2020). SARS-CoV-2 ORF9c Is a membrane-associated protein that suppresses antiviral responses in cells. bioRxiv 256776.

Angelini, M.M., Akhlaghpour, M., Neuman, B.W., and Buchmeier, M.J. (2013). Severe acute respiratory syndrome coronavirus nonstructural proteins 3, 4, and 6 induce double-membrane vesicles. mBio 4, e00524–13.

Banks, L., Pim, D., and Thomas, M. (2003). Viruses and the 26S proteasome: hacking into destruction.Trends Biochem. Sci. 28, 452–459.

Bennett, S.R., Carbone, F.R., Karamalis, F., Flavell, R.A., Miller, J.F., and Heath, W.R. (1998). Help for cytotoxic-T-cell responses is mediated by CD40 signalling. Nature 393, 478–480.

Blackburn, J.B., D’Souza, Z., and Lupashin, V.V. (2019). Maintaining order: COG complex controls golgi trafficking, processing, and sorting. FEBS Lett. 593, 2466–2487.

Blanco-Melo, D., Nilsson-Payant, B.E., Liu, W.-C., Uhl, S., Hoagland, D., Møller, R., Jordan, T.X., Oishi, K., Panis, M., Sachs, D., et al. (2020). Imbalanced host response to SARS-CoV-2 drives development of COVID-19. Cell 181, 1036–1045.e9.

Boehm, C.M., Obado, S., Gadelha, C., Kaupisch, A., Manna, P.T., Gould, G.W., Munson, M., Chait, B.T., Rout, M.P., and Field, M.C. (2017). The trypanosome exocyst: a conserved structure revealing a new role in endocytosis. PLoS Pathog. 13, e1006063.

Bouhaddou, M., Memon, D., Meyer, B., White, K.M., Rezelj, V.V., Correa Marrero, M., Polacco, B.J., Melnyk, J.E., Ulferts, S., Kaake, R.M., et al. (2020). The global phosphorylation landscape of SARS-CoV-2 infection. Cell 182, 685–712.e19.

Braun, P. (2013). Reproducibility restored--on toward the human interactome. Nat. Methods 10, 301, 303.

Braun, P., Tasan, M., Dreze, M., Barrios-Rodiles, M., Lemmens, I., Yu, H., Sahalie, J.M., Murray, R.R., Roncari, L., de Smet, A.-S., et al. (2009). An experimentally derived confidence score for binary protein-protein interactions. Nat. Methods 6, 91–97.

Calderwood, M.A., Venkatesan, K., Xing, L., Chase, M.R., Vazquez, A., Holthaus, A.M., Ewence, A.E., Li, N., Hirozane-Kishikawa, T., Hill, D.E., et al. (2007). Epstein-Barr virus and virus human protein interaction maps. Proc. Natl. Acad. Sci. U. S. A. 104, 7606–7611.

Carfì, A., Bernabei, R., Landi, F., and Gemelli Against COVID-19 Post-Acute Care Study Group (2020). Persistent symptoms in patients after acute COVID-19. JAMA 324, 603–605.

Cassel, J.A., and Reitz, A.B. (2013). Ubiquilin-2 (UBQLN2) binds with high affinity to the C-terminal region of TDP-43 and modulates TDP-43 levels in H4 cells: Characterization of inhibition by nucleic acids and 4-aminoquinolines. Biochimica et Biophysica Acta (BBA) - Proteins and Proteomics 1834, 964–971.

Chan, J.F.-W., Kok, K.-H., Zhu, Z., Chu, H., To, K.K.-W., Yuan, S., and Yuen, K.-Y. (2020). Genomic characterization of the 2019 novel human-pathogenic coronavirus isolated from a patient with atypical pneumonia after visiting Wuhan. Emerging Microbes & Infections 9, 221–236.

Choi, S.G., Olivet, J., Cassonnet, P., Vidalain, P.-O., Luck, K., Lambourne, L., Spirohn, K., Lemmens, I., Dos Santos, M., Demeret, C., et al. (2019). Maximizing binary interactome mapping with a minimal number of assays. Nature Communications 10, 3907.

Conibear, E., Cleck, J.N., and Stevens, T.H. (2003). Vps51p mediates the association of the GARP (Vps52/53/54) complex with the late Golgi t-SNARE Tlg1p. Mol. Biol. Cell 14, 1610–1623.

Conwell, S.E., White, A.E., Harper, J.W., and Knipe, D.M. (2015). Identification of TRIM27 as a novel degradation target of herpes simplex virus 1 ICP0. J. Virol. 89, 220–229.

Cusick, M.E., Yu, H., Smolyar, A., Venkatesan, K., Carvunis, A.-R., Simonis, N., Rual, J.-F., Borick, H., Braun, P., Dreze, M., et al. (2009). Literature-curated protein interaction datasets. Nat. Methods 6, 39–46.

Fajgenbaum, D.C., and June, C.H. (2020). Cytokine storm. N. Engl. J. Med. 383, 2255–2273.

Fanunza, E., Frau, A., Corona, A., and Tramontano, E. (2019). Insights into ebola virus VP35 and VP24 interferon inhibitory functions and their initial exploitation as drug targets. Infect. Disord. Drug Targets 19, 362–374.

Fields, S., and Song, O. (1989). A novel genetic system to detect protein-protein interactions. Nature 340, 245–246.

Fu, B., Wang, L., Ding, H., Schwamborn, J.C., Li, S., and Dorf, M.E. (2015). TRIM32 Senses and restricts influenza a virus by ubiquitination of PB1 polymerase. PLoS Pathog. 11, e1004960.

van Gent, M., Sparrer, K.M.J., and Gack, M.U. (2018). TRIM proteins and their roles in antiviral host defenses. Annu Rev Virol 5, 385–405.

Gilpin, K.M., Chang, L., and Monteiro, M.J. (2015). ALS-linked mutations in ubiquilin-2 or hnRNPA1 reduce interaction between ubiquilin-2 and hnRNPA1. Hum. Mol. Genet. 24, 2565–2577.

Gordon, D.E., Hiatt, J., Bouhaddou, M., Rezelj, V.V., Ulferts, S., Braberg, H., Jureka, A.S., Obernier, K., Guo, J.Z., Batra, J., et al. (2020b). Comparative host-coronavirus protein interaction networks reveal pan-viral disease mechanisms. Science 370, eabe9403.

Gordon, D.E., Jang, G.M., Bouhaddou, M., Xu, J., Obernier, K., White, K.M., O’Meara, M.J., Rezelj, V.V., Guo, J.Z., Swaney, D.L., et al. (2020a). A SARS-CoV-2 protein interaction map reveals targets for drug repurposing. Nature 583, 459–468.

Grubaugh, N.D., Hodcroft, E.B., Fauver, J.R., Phelan, A.L., and Cevik, M. (2021). Public health actions to control new SARS-CoV-2 variants. Cell 184, 1127–1132.

Gulbahce, N., Yan, H., Dricot, A., Padi, M., Byrdsong, D., Franchi, R., Lee, D.-S., Rozenblatt-Rosen, O., Mar, J.C., Calderwood, M.A., et al. (2012). Viral perturbations of host networks reflect disease etiology. PLoS Comput. Biol. 8, e1002531.

Gupta, N., Badeaux, M., Liu, Y., Naxerova, K., Sgroi, D., Munn, L.L., Jain, R.K., and Garkavtsev, I. (2017). Stress granule-associated protein G3BP2 regulates breast tumor initiation. Proc. Natl. Acad. Sci. U. S. A. 114, 1033–1038.

Hadjadj, J., Yatim, N., Barnabei, L., Corneau, A., Boussier, J., Smith, N., Péré, H., Charbit, B., Bondet, V., Chenevier-Gobeaux, C., et al. (2020). Impaired type I interferon activity and inflammatory responses in severe COVID-19 patients. Science 369, 718–724.

Hikmet, F., Méar, L., Edvinsson, Å., Micke, P., Uhlén, M., and Lindskog, C. (2020). The protein expression profile of ACE2 in human tissues. Mol. Syst. Biol. 16, e9610.

Huang, C., Wang, Y., Li, X., Ren, L., Zhao, J., Hu, Y., Zhang, L., Fan, G., Xu, J., Gu, X., et al. (2020). Clinical features of patients infected with 2019 novel coronavirus in Wuhan, China. Lancet 395, 497–506.

Jungreis, I., Nelson, C.W., Ardern, Z., Finkel, Y., Krogan, N.J., Sato, K., Ziebuhr, J., Stern-Ginossar, N., Pavesi, A., Firth, A.E., et al. (2020). Conflicting and ambiguous names of overlapping ORFs in SARS-CoV-2: A homology-based resolution. Preprints 2020120048.

Kanber, D., Giltay, J., Wieczorek, D., Zogel, C., Hochstenbach, R., Caliebe, A., Kuechler, A., Horsthemke, B., and Buiting, K. (2009). A paternal deletion of MKRN3, MAGEL2 and NDN does not result in Prader–Willi syndrome. Eur. J. Hum. Genet. 17, 582–590.

Karki, R., Sharma, B.R., Tuladhar, S., Williams, E.P., Zalduondo, L., Samir, P., Zheng, M., Sundaram, B., Banoth, B., Malireddi, R.K.S., et al. (2021). Synergism of TNF-α and IFN-γ triggers inflammatory cell death, tissue damage, and mortality in SARS-CoV-2 infection and cytokine shock syndromes. Cell 184, 149–168.e17.

Kim, D.-K., Knapp, J.J., Kuang, D., Chawla, A., Cassonnet, P., Lee, H., Sheykhkarimli, D., Samavarchi-Tehrani, P., Abdouni, H., Rayhan, A., et al. (2020). A comprehensive, flexible collection of SARS-CoV-2 coding regions. G3 10, 3399–3402.

Kim, D.-K., Sheykhkarimli, D., Kishore, N., Rayhan, A., Kuang, D., Li, R., Li, S., Škalič, M., Colobella, C., Cote, A.G., et al. (2021). Global dynamics of environment-dependent interactomes. Unpublished.

de Lange, T. (2005). Shelterin: the protein complex that shapes and safeguards human telomeres. Genes Dev. 19, 2100–2110.

Laurent, E.M.N., Sofianatos, Y., Komarova, A., Gimeno, J.-P., Tehrani, P.S., Kim, D.-K., Abdouni, H., Duhamel, M., Cassonnet, P., Knapp, J.J., et al. (2020). Global BioID-based SARS-CoV-2 proteins proximal interactome unveils novel ties between viral polypeptides and host factors involved in multiple COVID19-associated mechanisms. bioRxiv 272955.

Lee, B.O., Hartson, L., and Randall, T.D. (2003). CD40-deficient, influenza-specific CD8 memory T cells develop and function normally in a CD40-sufficient environment. J. Exp. Med. 198, 1759–1764.

Li, J., Guo, M., Tian, X., Wang, X., Yang, X., Wu, P., Liu, C., Xiao, Z., Qu, Y., Yin, Y., et al. (2021). Virus-host interactome and proteomic survey reveal potential virulence factors influencing SARS-CoV-2 pathogenesis. Med (N Y) 2, 99–112.e7.

Liu, Z., Miller, D., Li, F., Liu, X., and Levy, S.F. (2020). A large accessory protein interactome is rewired across environments. eLife 9, e62365.

Lu, H., Zhou, Q., He, J., Jiang, Z., Peng, C., Tong, R., and Shi, J. (2020). Recent advances in the development of protein-protein interactions modulators: mechanisms and clinical trials. Signal Transduct Target Ther 5, 213.

Luck, K., Kim, D.-K., Lambourne, L., Spirohn, K., Begg, B.E., Bian, W., Brignall, R., Cafarelli, T., Campos-Laborie, F.J., Charloteaux, B., et al. (2020). A reference map of the human binary protein interactome. Nature 580, 402–408.

Matthews, L.R., Vaglio, P., Reboul, J., Ge, H., Davis, B.P., Garrels, J., Vincent, S., and Vidal, M. (2001). Identification of potential interaction networks using sequence-based searches for conserved protein-protein interactions or “interologs.” Genome Res. 11, 2120–2126.

Miorin, L., Kehrer, T., Sanchez-Aparicio, M.T., Zhang, K., Cohen, P., Patel, R.S., Cupic, A., Makio, T., Mei, M., Moreno, E., et al. (2020). SARS-CoV-2 Orf6 hijacks Nup98 to block STAT nuclear import and antagonize interferon signaling. Proc. Natl. Acad. Sci. U. S. A. 117, 28344–28354.

Muller, M., Jacob, Y., Jones, L., Weiss, A., Brino, L., Chantier, T., Lotteau, V., Favre, M., and Demeret, C. (2012). Large scale genotype comparison of human papillomavirus E2-host interaction networks provides new insights for e2 molecular functions. PLoS Pathog. 8, e1002761.

Nabeel-Shah, S., Lee, H., Ahmed, N., and Marcon, E. (2020). SARS-CoV-2 Nucleocapsid protein attenuates stress granule formation and alters gene expression via direct interaction with host mRNAs. bioRxiv 342113.

Neveu, G., Cassonnet, P., Vidalain, P.-O., Rolloy, C., Mendoza, J., Jones, L., Tangy, F., Muller, M., Demeret, C., Tafforeau, L., et al. (2012). Comparative analysis of virus-host interactomes with a mammalian high-throughput protein complementation assay based on Gaussia princeps luciferase. Methods 58, 349–359.

Ochsenbein, A.F., Riddell, S.R., Brown, M., Corey, L., Baerlocher, G.M., Lansdorp, P.M., and Greenberg, P.D. (2004). CD27 expression promotes long-term survival of functional effector–memory CD8+cytotoxic T lymphocytes in HIV-infected patients. J. Exp. Med. 200, 1407–1417.

Orchard, S., Ammari, M., Aranda, B., Breuza, L., Briganti, L., Broackes-Carter, F., Campbell, N.H., Chavali, G., Chen, C., del-Toro, N., et al. (2014). The MIntAct project--IntAct as a common curation platform for 11 molecular interaction databases. Nucleic Acids Res. 42, D358–D363.

Peng, Z., Shi, T., and Ma, D. (2010). RNF122: a novel ubiquitin ligase associated with calcium-modulating cyclophilin ligand. BMC Cell Biol. 11, 41.

Pfefferle, S., Schöpf, J., Kögl, M., Friedel, C.C., Müller, M.A., Carbajo-Lozoya, J., Stellberger, T., von Dall’Armi, E., Herzog, P., Kallies, S., et al. (2011). The SARS-coronavirus-host interactome: identification of cyclophilins as target for pan-coronavirus inhibitors. PLoS Pathog. 7, e1002331.

Reineke, L.C., and Lloyd, R.E. (2015). The stress granule protein G3BP1 recruits protein kinase R to promote multiple innate immune antiviral responses. J. Virol. 89, 2575–2589.

Renaud, L., Picher-Martel, V., Codron, P., and Julien, J.-P. (2019). Key role of UBQLN2 in pathogenesis of amyotrophic lateral sclerosis and frontotemporal dementia. Acta Neuropathol Commun 7, 103.

Rolland, T., Taşan, M., Charloteaux, B., Pevzner, S.J., Zhong, Q., Sahni, N., Yi, S., Lemmens, I., Fontanillo, C., Mosca, R., et al. (2014). A proteome-scale map of the human interactome network. Cell 159, 1212–1226.

Rual, J.-F., Venkatesan, K., Hao, T., Hirozane-Kishikawa, T., Dricot, A., Li, N., Berriz, G.F., Gibbons, F.D., Dreze, M., Ayivi-Guedehoussou, N., et al. (2005). Towards a proteome-scale map of the human protein–protein interaction network. Nature 437, 1173–1178.

Salipante, S.J., Rojas, M.E.B., Korkmaz, B., Duan, Z., Wechsler, J., Benson, K.F., Person, R.E., Grimes, H.L., and Horwitz, M.S. (2009). Contributions to neutropenia from PFAAP5 (N4BP2L2), a novel protein mediating transcriptional repressor cooperation between Gfi1 and neutrophil elastase. Mol. Cell. Biol. 29, 4394–4405.

Samavarchi-Tehrani, P., Abdouni, H., Knight, J.D.R., Astori, A., Samson, R., Lin, Z.-Y., Kim, D.-K., Knapp, J.J., St-Germain, J., Go, C.D., et al. (2020). A SARS-CoV-2 – host proximity interactome. bioRxiv 282103.

Sicari, D., Chatziioannou, A., Koutsandreas, T., Sitia, R., and Chevet, E. (2020). Role of the early secretory pathway in SARS-CoV-2 infection. J. Cell Biol. 219.

St-Germain, J.R., Astori, A., Samavarchi-Tehrani, P., Abdouni, H., Macwan, V., Kim, D.-K., Knapp, J.J., Roth, F.P., Gingras, A.-C., and Raught, B. (2020). A SARS-CoV-2 BioID-based virus-host membrane protein interactome and virus peptide compendium: new proteomics resources for COVID-19 research. bioRxiv 269175.

Stukalov, A., Girault, V., Grass, V., Bergant, V., Karayel, O., Urban, C., Haas, D.A., Huang, Y., Oubraham, L., Wang, A., et al. (2020). Multi-level proteomics reveals host-perturbation strategies of SARS-CoV-2 and SARS-CoV. bioRxiv 156455.

Tang, Q., Wu, P., Chen, H., and Li, G. (2018). Pleiotropic roles of the ubiquitin-proteasome system during viral propagation. Life Sci. 207, 350–354.

Thoms, M., Buschauer, R., Ameismeier, M., Koepke, L., Denk, T., Hirschenberger, M., Kratzat, H., Hayn, M., Mackens-Kiani, T., Cheng, J., et al. (2020). Structural basis for translational shutdown and immune evasion by the Nsp1 protein of SARS-CoV-2. Science 369, 1249–1255.

Toes, R.E.M., Schoenberger, S.P., van der Voort, E.I.H., Offringa, R., and Melief, C.J.M. (1998). CD40–CD40 ligand interactions and their role in cytotoxic T lymphocyte priming and anti-tumor immunity. Semin. Immunol. 10, 443–448.

van de Ven, K., and Borst, J. (2015). Targeting the T-cell co-stimulatory CD27/CD70 pathway in cancer immunotherapy: rationale and potential. Immunotherapy 7, 655–667.

Walhout, A.J., Sordella, R., Lu, X., Hartley, J.L., Temple, G.F., Brasch, M.A., Thierry-Mieg, N., and Vidal, M. (2000). Protein interaction mapping in C. elegans using proteins involved in vulval development.Science 287, 116–122.

Wandinger-Ness, A., and Zerial, M. (2014). Rab proteins and the compartmentalization of the endosomal system. Cold Spring Harbor Perspectives in Biology 6, a022616–a022616.

Wang, Y., Sun, Y., Wu, A., Xu, S., Pan, R., Zeng, C., Jin, X., Ge, X., Shi, Z., Ahola, T., et al. (2015). Coronavirus nsp10/nsp16 methyltransferase can be targeted by nsp10-derived peptide in vitro and in vivo to reduce replication and pathogenesis. J. Virol. 89, 8416–8427.

Wehrens, E.J., Wong, K., Gupta, A., Benedict, C., and Zuniga, E. (2016). IL-27 suppresses CD4 and CD8 T cell cytotoxicity and viral control during cytomegalovirus infection. The Journal of Immunology 196, 217.4–217.4.

Wehrens, E.J., Wong, K.A., Gupta, A., Khan, A., Benedict, C.A., and Zuniga, E.I. (2018). IL-27 regulates the number, function and cytotoxic program of antiviral CD4 T cells and promotes cytomegalovirus persistence. PLoS One 13, e0201249.

Wu, A., Peng, Y., Huang, B., Ding, X., Wang, X., Niu, P., Meng, J., Zhu, Z., Zhang, Z., Wang, J., et al. (2020a). Genome composition and divergence of the novel coronavirus (2019-nCoV) originating in china. Cell Host & Microbe 27, 325–328.

Wu, F., Zhao, S., Yu, B., Chen, Y.-M., Wang, W., Song, Z.-G., Hu, Y., Tao, Z.-W., Tian, J.-H., Pei, Y.-Y., et al. (2020b). A new coronavirus associated with human respiratory disease in China. Nature 579, 265–269.

Yachie, N., Petsalaki, E., Mellor, J.C., Weile, J., Jacob, Y., Verby, M., Ozturk, S.B., Li, S., Cote, A.G., Mosca, R., et al. (2016). Pooled-matrix protein interaction screens using Barcode Fusion Genetics. Mol.Syst. Biol. 12, 863.

Yang, Q., Liu, T.-T., Lin, H., Zhang, M., Wei, J., Luo, W.-W., Hu, Y.-H., Zhong, B., Hu, M.-M., and Shu, H.-B. (2017). TRIM32-TAX1BP1-dependent selective autophagic degradation of TRIF negatively regulates TLR3/4-mediated innate immune responses. PLoS Pathog. 13, e1006600.

Yu, H., Luscombe, N.M., Lu, H.X., Zhu, X., Xia, Y., Han, J.-D.J., Bertin, N., Chung, S., Vidal, M., and Gerstein, M. (2004). Annotation transfer between genomes: protein-protein interologs and protein-DNA regulogs. Genome Res. 14, 1107–1118.

Yu, H., Braun, P., Yildirim, M.A., Lemmens, I., Venkatesan, K., Sahalie, J., Hirozane-Kishikawa, T., Gebreab, F., Li, N., Simonis, N., et al. (2008). High-quality binary protein interaction map of the yeast interactome network. Science 322, 104–110.

Yuen, C.-K., Lam, J.-Y., Wong, W.-M., Mak, L.-F., Wang, X., Chu, H., Cai, J.-P., Jin, D.-Y., To, K.K.-W., Chan, J.F.-W., et al. (2020). SARS-CoV-2 nsp13, nsp14, nsp15 and orf6 function as potent interferon antagonists. Emerg. Microbes Infect. 9, 1418–1428.

Zarrinpar, A., Park, S.-H., and Lim, W.A. (2003). Optimization of specificity in a cellular protein interaction network by negative selection. Nature 426, 676–680.

Zha, J., Han, K.-J., Xu, L.-G., He, W., Zhou, Q., Chen, D., Zhai, Z., and Shu, H.-B. (2006). The Ret finger protein inhibits signaling mediated by the noncanonical and canonical IkappaB kinase family members.J. Immunol. 176, 1072–1080.

Zhang, J., Hu, M.-M., Wang, Y.-Y., and Shu, H.-B. (2012). TRIM32 protein modulates type I interferon induction and cellular antiviral response by targeting MITA/STING protein for K63-linked ubiquitination. J.Biol. Chem. 287, 28646–28655.

Zhang, R., Poustovoitov, M.V., Ye, X., Santos, H.A., Chen, W., Daganzo, S.M., Erzberger, J.P., Serebriiskii, I.G., Canutescu, A.A., Dunbrack, R.L., et al. (2005). Formation of MacroH2A-containing senescence-associated heterochromatin foci and senescence driven by ASF1a and HIRA. Dev. Cell 8, 19–30.

Zheng, M., Gao, Y., Wang, G., Song, G., Liu, S., Sun, D., Xu, Y., and Tian, Z. (2020). Functional exhaustion of antiviral lymphocytes in COVID-19 patients. Cell. Mol. Immunol. 17, 533–535.

Zheng, Q., Hou, J., Zhou, Y., Yang, Y., Xie, B., and Cao, X. (2015). Siglec1 suppresses antiviral innate immune response by inducing TBK1 degradation via the ubiquitin ligase TRIM27. Cell Res. 25, 1121–1136.

Zhong, Q., Pevzner, S.J., Hao, T., Wang, Y., Mosca, R., Menche, J., Taipale, M., Taşan, M., Fan, C., Yang, X., et al. (2016). An inter-species protein-protein interaction network across vast evolutionary distance. Mol. Syst. Biol. 12, 865.

