## Supplementary material for "A map of binary SARS-CoV-2 protein interactions implicates host immune regulation and ubiquitination": Method

### STAR METHOD

#### KEY RESOURCES TABLE

| REAGENT or RESOURCE | SOURCE | IDENTIFIER |
|---|---|---|
| **Chemicals** | | |
| carbenicillin | Bioshop | CAR544.10 |
| chloramphenicol | Bioshop | CLR201.10 |
| Omni plates | ThermoFisher Scientific | 12-565-296 |
| 3-amino-1,2,4-triazole (3-AT) | Sigma Aldrich | A8056 |
| cycloheximide (CYH) | Sigma Aldrich | C7698 |
| Phosphate-Buffered Saline (PBS) | Wisent | 311-425-CL |
| Propidium iodide (PI) | Bioship | PPI888.10 |
| **Critical Commercial Assays** | | |
| HiPure Plasmid Filter Maxiprep Kit | Invitrogen | K210016 |
| Gateway LR reactions | ThermoFisher Scientific | 11791100 |
| Yeast Plasmid Miniprep Kits | Zymoprep | D2004 |
| NucleoSpin Gel and PCR Clean-up kit | Macherey-Nagel | 740609.250 |
| Qubit | Invitrogen | Q32851 |
| Agencourt AMPure XP | Beckman Coulter | A63881 |
| KAPA library quantification kit | KAPA Biosystems | KK4824 |
| KAPA SYBR FAST qPCR Master Mix | KAPA Biosystems | KM4101 |
| 96-well DNA extraction kit | Qiagen | 27193 |
| **Deposited Data** | | |
| SARS-CoV-2 Genomes (Wuhan/IVDC-HB-01/2019 (HB01), Wuhan/IVDCHB-04/2019 (HB04) and Wuhan/IVDC-HB-05/2019 (HB05)) | (Wu et al., 2020b) | NC_045512 |
| Human ORFeome collection v9.1 | (Luck et al., 2020; The | N/A |

| | | |
|---|---|---|
| | ORFeome Collaboration, 2016) | |
| barcoded human ORFeome (bhORFeome) | (Luck et al., 2020) | N/A |
| hsPRS-v2 (Human Positive Reference Set, v2) | (Braun et al., 2009; Choi et al., 2019) | N/A |
| hsRRS-v2 (Human Random Reference Set, v2) | (Braun et al., 2009; Choi et al., 2019) | N/A |
| virus-host literature binary multiple reference set (vhLit-BM) | This manuscript | N/A |
| viral-host Random Reference Set (vhRRS) | This manuscript | N/A |
| **Experimental Models: Cell lines** | | |
| ccdB-resistant electrocompetent cells | (Yachie et al., 2016) | N/A |
| One Shot ccdB Survival 2 T1R Competent Cells | Invitrogen | A10460 |
| Yeast (Y8800, MATa, Prey strains) | (Altmann et al., 2018, 2020; Choi et al., 2019) | N/A |
| Yeast (Y8930, MATα, Bait strains) | (Altmann et al., 2018, 2020; Choi et al., 2019) | N/A |
| Yeast (RY3011, MATa, Prey strains) | (Kim et al., 2021) | N/A |
| Yeast (RY3031, MATα, Bait strains) | (Kim et al., 2021) | N/A |
| **Experimental Models: Organisms/Strains** | | |
| pENTR223.1*Sfi | Invitrogen | N/A |
| pAR068 (C-terminal AD fusion, 2μ origin) | (Yachie et al., 2016) | N/A |
| pHiDEST-AD (N-terminal AD fusion, CEN origin) | (Yachie et al., 2016) | N/A |
| pHiDEST-DB (N-terminal DB fusion, CEN origin) | (Yachie et al., 2016) | N/A |
| pPC86 (N-terminal AD fusion, CEN origin) | (Altmann et al., 2018, 2020) | N/A |
| pDEST-N2H (-N1, -N2, -C1 and -C2) | (Choi et al., 2019) | N/A |
| **Software and Algorithms** | | |
| bowtie2 (v2.2.3) | (Langmead and Salzberg, 2012) | RRID:SCR_016368 (http://bowtie-bio.sourcefo |

| | | |
|---|---|---|
| | | rge.net/bowtie2/index.shtml) |
| bcl2fastq2 (v2.20.0.422) | Illumina | N/A |
| Cytoscape (v3.8.2) | (Shannon et al., 2003) | N/A |
| Metascape web-platform | (Zhou et al., 2019) | N/A |
| **Others** | | |
| BioMatrix Robot | S&P Robotics | BM3-BC |

#### LEAD CONTACT AND MATERIALS AVAILABILITY

#### EXPERIMENTAL MODEL AND SUBJECT DETAILS

Yeast strains are listed in the Key Resources Table.

#### METHOD DETAILS

##### Selection & clone preparation of SARS-CoV-2 functional ORFs

*Gene synthesis of SARS-CoV-2 ORFs into Gateway compatible vectors*

Two independent collections of Gateway Entry plasmids containing the ORFs of SARS-CoV-2 were constructed, one for $HuSCI_{HIS3}$ and another for $HuSCI_{GFP}$.

The $HuSCI_{GFP}$ collection is described in our recent paper (Kim et al., 2020). Briefly, it includes all but one (NSP11, which is too short for Gateway cloning) of the codon-optimized ORFs of SARS-CoV-2, synthesized based on a published genome (Wu et al., 2020b), and then transferred to Gateway-compatible Entry plasmids, in both ‘open’ (without a stop codon, to enable C-terminal fusions) and ‘closed’ (with a stop codon) configurations. In a similar fashion, the $HuSCI_{HIS3}$ collection of Gateway-compatible Entry plasmids was based on SARS-CoV-2 ORFs from published viral genome sequences Wuhan/IVDC-HB-01/2019 (HB01), Wuhan/IVDCHB-04/2019 (HB04) and Wuhan/IVDC-HB-05/2019 (HB05) (Wu et al., 2020a). All viral ORFs for $HuSCI_{HIS3}$ were synthesized by Twist Bioscience, and include two linkers at the 5´ and 3´ends, respectively, which contain SfiI restriction sites for cloning into pENTR223.1*SfiI. Additionally, the 5´ linker contains an alternative translational start ATG flanked by BamHI sites, and the 3´ linker contains a stop codon flanked by PacI and AsiSI restriction sites, to replace the native translational stop. This design allows for the removal of either the start or stop codon, to accomodate in-frame fusion to N-terminal or C-terminal protein tags, respectively. For $HuSCI_{HIS3}$, synthesized viral ORFs were cloned into pENTR223.1 by SfiI restriction digest. The start ATG was then removed by BamHI digest and subsequent re-ligation.

In total, 28 viral ORFs were synthesized for $HuSCI_{GFP}$ and 27 for $HuSCI_{HIS3}$. Together, these included: NSP1-16 (except NSP11), all derived from ORF1ab; the four structural proteins S, E, M and N; and the 10 accessory ORFs ORF3A, 3B, 3D, 6, 7A, 7B, 8, 9B, 9C and 10 (Wu et al., 2020a)(**Supplementary Table 1A**).

*Preparation of barcoded Y2H destination plasmids*

Barcoded 'prey' (pAR068: C-terminal AD fusion, 2µ origin / pHiDEST-AD: N-terminal AD fusion, CEN origin), and 'bait' (pHiDEST-DB: N-terminal DB fusion, CEN origin) Destination plasmid vectors were prepared using our published protocols (Yachie et al., 2016), with the integration of the barcode locus at the SacI restriction site. Following Gibson assembly (Gibson et al., 2009), DNA was transformed (BioRad MicroPulser electroporator, 1652100) into *ccdB*-resistant electrocompetent cells, made from One Shot *ccdB* Survival 2 T1R Competent Cells (Invitrogen, A10460). After selection on Luria-Bertani (LB) agar plates with 100 µg/mL carbenicillin (Bioshop, CAR544.10) and 35 µg/mL chloramphenicol (Bioshop, CLR201.10), bacterial colonies, which are representative of clonal barcoded plasmids, were pooled and mixed thoroughly to generate a high complexity pool. Pooled barcoded 'bait' and 'prey' destination plasmid DNA were then isolated by maxiprep (Invitrogen, K210016).

*Isolation and sequencing of clonal barcoded null plasmids*

To isolate clonal barcoded plasmids, the destination plasmid pool was transformed into One Shot *ccdB* Survival 2 T1R Competent Cells (Invitrogen, A10460). Cells were selected on Omni plates (Fisher Scientific, 12-565-296) containing LB agar supplemented with 100 µg/mL carbenicillin and 35 µg/mL chloramphenicol, and incubated overnight at 37°C such that each plate yielded 500-1,500 isolated single colonies. Single colonies were picked using a Biomatrix robot (S&P Robotics) and arrayed into 384-well cell culture plates containing 80 µL of LB with carbenicillin (100 µg/mL) and chloramphenicol (35 µg/mL) in each well. The cell culture plates were sealed by foil seal (VWR, 60941-126) and incubated at 37°C for 16h.

To determine the sequence of each plasmid barcode, the Kiloseq procedure was performed as described (Weile et al., 2017) with modifications reflecting that we only used this approach to sequence barcodes for these empty vectors. In the first step, we performed colony PCR in 384-well plate format, in which well-specific indexing primers were attached to the barcode sequence. Amplified DNA was pooled plate-wise and DNA pools were purified with a PCR clean-up kit (Macherey-Nagel). Each plate-pool of DNA was then re-amplified with primers designed to add Illumina adapters in order to generate a library of amplicons such that each amplicon carried plate- and well-specific information, allowing multiple plates of amplicons to be sequenced together. Indexed barcoded libraries were assessed with the KAPA library quantification kit (Kapa Biosystems, KK4824) and sequenced using an Illumina NextSeq 500. Sequence analysis was carried out as described previously (Luck et al., 2020; Weile et al., 2017; Yachie et al., 2016).

*Subcloning of SARS-CoV-2 ORFs into Y2H Destination plasmids*

In preparation for the $HuSCI_{HIS3}$ Y2H screen, each viral ORF was moved to the Destination vectors pPC86 (N-terminal AD fusion, CEN origin) (Altmann et al., 2018, 2020) and pHiDEST-DB (N-terminal DB fusion, CEN origin) (Yachie et al., 2016) by Gateway recombinational cloning to produce AD-Y and DB-X N-terminal fusion constructs, respectively. Because viral ORF sequences had been synthesized and their sequences verified prior to cloning, successful recombination into destination plasmids was confirmed by PCR evaluation of insert size. The viral ORF sequences for all interactions identified in $HuSCI_{HIS3}$ were re-confirmed by Sanger sequencing post-screen (see below).

Gateway LR reactions (Gateway™ LR Clonase™ II Enzyme mix, 11791100) were performed for the entire $HuSCI_{GFP}$ library of SARS-CoV-2 ORFs, and the human gene ACE2 into three different configurations of barcoded destination plasmids (Luck et al., 2020; Yachie et al., 2016): pAR068 (C-terminal AD fusion, 2µ origin (high copy number)), pHiDEST-AD (N-terminal AD fusion, CEN origin (low copy number)), pHiDEST-DB (N-terminal DB fusion, CEN origin (low copy number)) such that each ORF was linked to two to six barcodes in every configuration. Gateway LR reactions were performed individually to ensure that each ORF was linked with known barcode sequences in the constructed destination plasmid collection. These collections of barcoded destination vectors were then sequence-verified to confirm ORF-barcode pairs using Sanger sequencing (TCAG, SickKids Hospital).

ORFs and associated barcode sequences are described in **Supplementary Table 3**.

**HIS3 based screening to generate the HuSCI$_{HIS3}$ map**

The Y2H$_{HIS3}$ screening pipeline is essentially as described (Altmann et al., 2018). Briefly, AD-Y and DB-X plasmids containing 27 viral ORFs were transformed into yeast strains Y8800 (MATa) and Y8930 (MATα), respectively. Individual DB-X viral fusion constructs were first mated with yeast containing AD-empty plasmid, in order to identify 'auto-activating' 'baits'. Only NSP1 showed a high level of activation of the *LYS2::GAL1-HIS3* reporter on selective plates containing synthetic complete media lacking leucine, tryptophan and histidine (SC-Leu-Trp-His), supplemented with 20 mM of 3-amino-1,2,4-triazole (3-AT) (Sigma Aldrich, A8056) and was therefore excluded from screening in this orientation. All 26 remaining DB-X viral ORFs were individually mated with 99 pools, each containing ~188 AD-Y human ORF constructs, obtained from the human ORFeome collection v9.1 from the Center for Cancer Systems biology (CCSB), comprising a total of 17,472 ORFs (Luck et al., 2020; The ORFeome Collaboration, 2016). For the reverse screening orientation, yeast cultures containing 27 AD-Y viral fusion constructs were pooled and the pool was mated with DB-X human fusion constructs individually, also derived from the human ORFeome collection v9.1. DB-X auto-activating constructs had been removed from the collection prior to screening, as previously described (Luck et al., 2020). Screening in both configurations was performed two times in order to increase sampling sensitivity.

Mating was performed by spotting 5 µL of each of the saturated haploid yeast cultures onto plates containing yeast extract peptone dextrose (YEPD) medium and incubating for 24 hours at 30°C. Yeast spots were then transferred onto SC-Leu-Trp-His + 1mM 3-AT plates using a replica plating block (Altmann et al., 2018, 2020) and incubated for 72h at 30°C.

Y2H-positive colonies were identified by growth on plates containing SC-Leu-Trp-His + 1mM 3-AT and 3 colonies per spot were picked and individually cultured in SC-Leu-Trp liquid medium and grown for 2 days at 30°C. This culture was spotted (5 µL) on SC-Leu-Trp plates and grown for another 2 days at 30°C. Yeast spots were then transferred onto SC-Leu-Trp-His + 1mM 3-AT and SC-Leu-His + 1mM 3-AT + 1mg/L (~3.4µM) cycloheximide (CYH) (Sigma Aldrich, C7698) plates using a replica plating block to test for cycloheximide-sensitive expression of the *LYS2::GAL1-HIS3* reporter, which enables identification of spontaneous DB-X auto-activators that can arise during screening (Yu et al., 2008). Positive scoring colonies (growth on SC-Leu-Trp-His + 3-AT, but not on SC-Leu-His + 3-AT + CYH) were picked and interaction candidates of AD-Y or DB-X human ORFs, respectively, were identified using end-read Sanger sequencing of PCR products (Altmann et al., 2018). Yeast strains corresponding to the identified AD-Y and DB-X human interaction candidates were picked from archival glycerol stocks, cultured in liquid medium and mated (as described above) one-by-one with all 27 DB-X or AD-Y viral ORFs, respectively. This binary verification mating was performed four times for the orientation of viral DB-X versus human AD-Y and Y2H interactions were scored positive if at least three of four repeats showed robust growth on SC-Leu-Trp-His + 1mM 3-AT plates. For viral AD-Y versus human DB-X, binary verification mating was done in triplicate and positive interactions scored if growth was observed in at least two out of three repeats.

In order to eliminate any operator bias when scoring yeast growth for positive interactions, yeast growth was scored automatically with a custom dilated convolutional neural network (Yu and Koltun, 2015). For training, previous datasets of more than 1,500 images of biochemically and functionally validated binary Y2H studies were used (Altmann et al., 2020). Each image was scaled to achieve equal pixel-distance between the yeast spots of different images. The images were then cropped and sliced, so that each yeast spot became a separate file. In addition, the mean grayscale image of all 96 yeast spots of the same plate was calculated. With this dataset, a simple front-end prediction module was trained. The front-end module consisted of six dilated convolutional layers with exponential increasing dilation rate and two dense layers at the end. After each layer except the last, a Leaky-ReLU activation was added (Maas et al., 2013). The model was optimized with a combination of Softmax and Cross entropy and an Adam Optimizer (Kingma and Ba, 2014). The model achieved an accuracy of >0.9 during all folds of a ten-fold cross-fold validation. All positive scores were confirmed by a trained researcher. One representative colony of all positive scoring pairs was picked from selective plates and the identities of DB-X and AD-Y were confirmed by Sanger sequencing of PCR products (Altmann et al., 2018).

#### GFP based screening to generate the HuSCI$_{GFP}$ map

*Generation of Y2H$_{GFP}$ strains for HuSCI$_{GFP}$*

Development of GFP-based Y2H technology, as well as generation of toolkit strains is as described in our forthcoming publication (Kim et al., 2021). Briefly, Y2H strains carrying multiple copies of GFP based Y2H reporters were engineered and empirically benchmarked with previously established PRS and RRS (hsPRS-v2 and hsRRS-v2) sets before proceeding with the experiments described in this study. Genotypes of toolkit strains are described in **Supplementary Table 4**.

*Generation of the barcoded human ORFeome (bhORFeome) for screening*

A Gateway Entry clone collection of the human ORFeome was available from CCSB (Luck et al., 2020; The ORFeome Collaboration, 2016). Based on this collection, *en masse* Gateway LR reactions using a pool of barcoded destination plasmids, bacterial transformations, colony picking and kiloseq were performed as described (Weile et al., 2017; Yachie et al., 2016). After obtaining the raw sequencing results, bowtie2 (v2.2.3) (Langmead and Salzberg, 2012) was used to align the reads in each well to the reference sequences. To optimize this collection of barcoded ORFs, this process was repeated three additional times, using only the missing ORFs from the previous rounds in subsequent barcoding attempts. The final clones were arrayed using an automated robot system (S&P Robotics Inc., BM3-BC). Approximately 1% of the clones were validated by Sanger sequencing with >95% success rate for both ORF and barcode sequences. Our final union of barcoded 'bait' (DB) and 'prey' (AD) plasmids, called 'barcoded human ORFeome (bhORFeome)', includes 16,747 fully sequence verified human ORFs (96% of the entire human ORFeome collection) with most (~95%) ORFs represented by 2 unique barcodes. Prior to yeast transformation of the bhORFeome, we cataloged the barcoded 'bait' and 'prey' destination plasmid collection into a 10-by-10 screening matrix consisting of ten DB and ten AD groups, respectively. Each haploid AD and DB group contains ~1,400 ORFs with two distinct sets of unique barcodes, and ~200 ORFs with a single unique barcode set.

*Transformation of SARS-CoV-2 destination plasmids into GFP based Y2H strains*

Each uniquely barcoded destination plasmid containing a sequence-verified SARS-CoV-2 ORF was transformed to the corresponding yeast strain (RY3011 for AD fusions, and RY3031 for DB fusions), individually in 96-well plates. After the transformants were selected by growth on SC media lacking specific amino acids (leucine for DB, and tryptophan for AD), 10 replica plates were made for each of the destination vector version collections. These plates were then scraped and pooled together to make glycerol stocks that are pools of all the barcoded SARS-CoV-2 ORFs plus the human ORF ACE2 in each plasmid configuration, to be screened using $Y2H_{GFP}$.

Pooled Glycerol stocks of haploid yeast:

| **Yeast Strain** | **Plasmid backbone** | **ORFs** | **Barcodes** |
|---|---|---|---|
| RY3031 | pHiDEST-DB<br>(N-term, DB, CEN) | SARS-CoV-2 + ACE2 | >= 2 each |
| RY3011 | pHiDEST-AD | SARS-CoV-2 + ACE2 | >= 2 each |

| | (N-term, AD, CEN) |
|---|---|

*Mating of pooled haploid yeast*

For $HuSCI_{GFP}$, multiple pooled matings were performed using the frozen haploid glycerol stock pools. Each of the 10 pools of human ORFs (in C-terminal AD fusion plasmids with 2μ origin; pAR068) were separately mixed with the pool of SARS-CoV-2 ORFs plus human ACE2 (in N-terminal DB fusion plasmids with CEN origins; pHiDEST-DB) for a total of 10 pooled matings to cover virus-host interactions. Another separate mating was done between the SARS-CoV-2 pools in both AD and DB fusion, CEN origin plasmids (pHiDEST-AD, pHiDEST-DB). These 11 pool-pool matings were set up to achieve >100x coverage with respect to the number of possible barcode combinations, while taking into account the number of viable colony-forming units (vCFUs) of the haploid glycerol stocks, and the expected mating efficiency. Negative controls consisting of empty destination vectors with matching configurations and known barcodes were also added to each mating. Matings were achieved by mixing together equal amounts of each haploid strain (plus added negative controls), spreading the cell mixture onto a double-strength YEPD plus adenine (2x YPAD) agar plate, and incubating at 30°C for 24h. Mating plates were then scraped with distilled water, and each mated culture was spread across 20 large (15 cm) SC-Leu-Trp petri plates, supplemented with additional histidine (80 mM) to ensure more equal representation of all strains (SC-Leu-Trp+10xHis), which are then incubated at 30°C for a further 72h. These plates were then scraped and assay-ready pooled diploid glycerol stocks were made for each of the 11 groups.

*Selection of yeast with interacting pair of DB-X and AD-Y by FACS*

Diploid pooled glycerol stocks were thawed on ice and inoculated into 1L flasks with a total starting viable CFU of 30M (to achieve ~100x coverage with respect to ORF pairs), and incubated at 30°C at 200 rpm for 24h. Smaller cultures (10 mL) of negative controls were also prepared. Before FACS, 'presort' cultures were prepared for each sample (2 x 10 mL of cultures with $OD_{600}$ 10) with doxycycline added (10 μg/mL) to these cultures to induce barcode swapping while these cultures incubate at 30°C for 24h (Yachie et al., 2016). To prepare for FACS, cells were concentrated by centrifugation (500 x g, 5min) and resuspended in Phosphate-Buffered Saline (PBS, Wisent, 311-425-CL) to a final $OD_{600}$ value of 10 to limit further growth. Negative controls were also centrifuged and resuspended in a final volume of 1mL PBS. Propidium iodide (PI, Bioshop, PPI888.10) was added to all samples and controls to a final concentration of 4 mg/L to discriminate dead yeast cells during FACS.

Using a mated negative control, the FACS gate for GFP positive cells was set to capture 0.1% of GFP-negative cells to yield a 0.01% false positive rate (FPR). FACS was performed on a Sony MA900 cell sorter with the aid of a core facility at Lunenfeld-Tanenbaum Research Institute (LTRI) at Mount Sinai Hospital in Toronto. We sorted over 100M cells per group for >100x coverage with respect to the number of ORF pairs.

After FACS, GFP positive cells for each sample were spread across 10 plates of selective agar media (SC-Leu-Trp+Ade+10xHis) and incubated at 30°C for 72h. These plates were then scraped, centrifuged, and resuspended into 2 x 10mL cultures at $OD_{600}$ of 10. Doxycycline (10 μg/mL) was then added to these cultures to induce barcode swapping while they were incubated at 30°C for 24h.

After doxycycline induction of both presort and GFP-positive samples, cells were harvested and plasmid DNA was extracted via yeast miniprep (Zymo Research, D2004). Fused barcodes were then amplified with primers that attach modified Illumina i5 and i7 adapters in order to uniquely identify each different sample. PCR products were run on a gel and the bright band at ~350 bp was size-selected and purified using the NucleoSpin Gel and PCR Clean-up kit (Macherey-Nagel, 740609.250).

*NGS library preparation and sequencing*

DNA concentrations were measured for each sample using a Qubit (Invitrogen, Q32851) and, using the DNA concentrations as a guide, samples were pooled together to ensure relative sequencing depth of each library was proportional to the number of protein pairs tested. The pooled NGS library was then processed with Agencourt AMPure XP (Beckman Coulter, A63881) to remove primer dimers, and quantified by qPCR (KAPA SYBR FAST qPCR Master Mix, KAPA Biosystems, KM4101). The library was sequenced on an Illumina NextSeq using a mid- or high-output 150 cycle kit to achieve the appropriate read-depth necessary for subsequent analysis.

*Read counting based on the expected barcodes*

The sequencing data was demultiplexed using bcl2fastq2 (v2.20.0.422) provided by Illumina with the following command:

“bcl2fastq -r 10 -p 20 -w 10 --no-lane-splitting --barcode-mismatches 1 --adapter-stringency 0.7 --ignore-missing-bcls --ignore-missing-filter --ignore-missing-positions”

After demultiplexing, the fastq files were aligned to the group specific reference files using bowtie2 (Langmead and Salzberg, 2012) with the following parameters:

For read 1: -q --norc --local --very-sensitive-local -t -p 23 --reorder

For read 2: -q --nofw --local --very-sensitive-local -t -p 23 --reorder

The reference files contain the expected barcode sequences for the ORFs in each group. After alignments, we count the number of reads mapped to each up/dn tag sequence. The reads were filtered out if the MAPQ score was lower than 20. Based on the BFG barcode recombination process (Yachie et al., 2016), paired end reads should map to up-up or dn-dn when an interaction is present. Hence, for each group, we build a matrix with DB-X and AD-Y as row names and virus ORFs as column names. We count the number of reads mapped to up-up and dn-dn separately and merge them together as our final read counts. The pipeline was implemented in Python2.7 and the source code can be found on Github (https://github.com/RyogaLi/BFG_Y2H/).

*__Calculation of interaction score and statistical test to identify confident PPI between SARS-CoV-2 and human ORFs__*

As described in (Yachie et al., 2016), we used the product of marginal frequencies of bait and prey strains to estimate the abundance of each diploid bait-prey strain in the pre-sort condition (“PreSort”). Interaction score (IS) was defined by

$$IS_{ij} = \frac{f_{ij}^{GFP}}{f_{ij}^{PreSort}}$$

$$f_i^{PreSort} = \sum_i c_{ij}^{PreSort} / \sum_j [\sum_i c_{ij}^{PreSort}] \quad f_j^{PreSort} = \sum_j c_{ij}^{PreSort} / \sum_i [\sum_j c_{ij}^{PreSort}]$$

$$f_{ij}^{PreSort} = \max(f_i^{PreSort}, f_{AD}^{Floor}) \times \max(f_j^{PreSort}, f_{DB}^{Floor}) \quad f_{AD}^{Floor} = 10^{-5} \quad f_{DB}^{Floor} = 10^{-4}$$

$$f_{ij}^{GFP} = c_{ij}^{GFP} / \sum_{ij} c_{ij}^{GFP}$$

with c, read count; i, AD barcode; j, DB barcode; f. frequency.

For each DB barcode, we identified the score threshold achieving a 1% false positive rate (FPR) using the 960 AD null barcodes included in this screen. An interaction was accepted as positive only if the ORF pair was above this IS score threshold for ≥ 2 barcode pairs.

*__Calculation of interaction score to identify confident PPI between SARS-CoV-2 ORFs__*

For intraviral screening, adding in the 100 AD-null controls would have enlarged the screening space by a factor of >100 over the actual protein pairs being screened. For this reason, we did not add the negative controls in this screen to use as a 1% false positive rate. Instead, we used a similar method to the one that was used for viral-human PPIs: we only accepted as interactions those protein pairs for which the frequency of barcode pairs was 1000 times greater than the median frequency of the corresponding each DB barcode for ≥3 independent barcode pairs, similar to the scoring method previously used for BFG-Y2H with HIS3-based growth selection (Yachie et al., 2016).

*Pairwise retesting using HIS3 selection*

Candidate interaction pairs for $HuSCI_{GFP}$ were retested in a pairwise HIS3 growth-based Y2H assay to verify the initial interaction result, following the same protocol as outlined above for the $Y2H_{HIS3}$ screen. Briefly, identified AD-Y human and DB-X viral interaction candidates were picked from archival glycerol stocks, cultured in liquid medium and mated (as described above) one-by-one with barcode replicates pooled together prior to mating. DB-X viral interaction candidates were also individually mated with AD-empty controls to assess whether they are classed as 'auto-activators'. Viral ORFs NSP1 and NSP12 were subsequently omitted from this retesting due to DB auto-activation. Interactions were scored after growing these individually mated interacting pairs on selection medium lacking histidine (SC-Leu-Trp-His), and medium lacking histidine supplemented with 3-AT (SC-Leu-Trp-His+1mM 3-AT) for stronger interactions. After 72-96h of yeast growth these pairwise tests were scored according to the standardized scoring method used for the $Y2H_{HIS3}$ screen (Altmann et al., 2018, 2020). Any interacting pairs that scored above the threshold for positive interactions (score ≥ 3) were collected together and subsequently sent for validation.

*Benchmarking $Y2H_{HIS3}$ and $Y2H_{GFP}$ against positive and negative control sets*

$Y2H_{HIS3}$ toolkit strains were benchmarked previously (Braun et al., 2009). Briefly, bait and prey constructs constituting 92 hsPRSv1 and 92 hsRRSv1 pairs were tested using the identical protocol to the one used to verify $HuSCI_{HIS3}$. $Y2H_{HIS3}$ recovered 23 out of 92 hsPRS-v1 pairs when tested in both bait-prey configurations corresponding to an assay sensitivity of $S_{a\text{-}HIS3}$ = 25%. A different version of $Y2H_{GFP}$ using low copy plasmids and N-terminally fused hybrid proteins ($lcnY2H_{GFP}$) was benchmarked using 84 pairs of hsPRS-v1 and 92 pairs of hsRRS-v1 sets and flow cytometry was used to score for interactions based on percentage of singlets in GFP-positive gate, which was set using empty bait and prey constructs. In addition, $lcnY2H_{GFP}$ was benchmarked in a pooled setting using all possible combinations of proteins constituting 78 hsPRS-v2 and 77 hsRRS-v2 pairs supplemented with a 14 pairs of Y2H-positive controls defined as Calibration Set (CS), (Yachie et al., 2016). The experiment was carried out and interactions were scored as described above, except that no empirical null distribution was used.

Assay sensitivity can be defined as a fraction of true interactions that can be detected by a given assay. Sampling sensitivity can be defined as a fraction of detectable true interactions that can be recovered by the screen. Let assay sensitivity be $S_a$ and sampling sensitivity be $S_s$, such that sensitivity of a given screen S can be calculated as $S = S_a \times S_s$. When sampling is saturated by repeated screens ($S_s = 1$), the screening sensitivity is equal to assay sensitivity, *i.e.* $S = S_a$. In pairwise settings $S_s = 1$ and the assay sensitivity is given by the fraction of hsPRSv1/v2 pairs that score positive. Sampling sensitivity of $Y2H_{HIS3}$ has previously shown to be $S_{s\text{-}HIS3} = 60\%$ (Altmann et al., 2018) after two repeats, yielding screening sensitivity of $S_{HIS3} = S_{a\text{-}HIS3} \times S_{s\text{-}HIS3} = 0.25 \times 0.6 = 15\%$ . Given that $Y2H_{HIS3}$ screen had a search space completeness of 83% ($T_{HIS3}$=83%), the completion level of the map generated by $Y2H_{HIS3}$ is $C_{HIS3} = T_{HIS3} \times S_{HIS3} = 0.83 \times 0.15 = 12.45\%$. lcn$Y2H_{GFP}$ recovered 12 out of 82 ($S_{a\text{-}lcnGFP} = 15\%$) hsPRSv1 pairs when tested in a pairwise single bait-prey configuration and 8 of 92 (9%, $S_{s\text{-}lcnGFP} = 9/15 = 60\%$) hsPRSv2+CS pairs when tested in a pooled single bait-prey configuration, yielding $S_{lcnGFP} = S_{a\text{-}lcnGFP} \times S_{s\text{-}lcnGFP} = 0.15 \times 0.6 = 9\%$. It has been previously shown that using high-copy C-terminal fusions increases sensitivity by ~50% without affecting the precision (Luck et al., 2020). Thus, screening sensitivity of $Y2H_{GFP}$ was modeled from that of lcn$Y2H_{GFP}$ as $S_{GFP} = S_{lcnGFP} \times 1.5 = 9\% * 1.5 = 13.5\%$. Given that $Y2H_{GFP}$ covered 70% ($T_{GFP}$=70%) of all possible virus-human protein combinations, the completion level of the $Y2H_{GFP}$ dataset is $C_{GFP} = T_{GFP} \times S_{GFP} = 0.70 \times 0.135 = 9.45\%$. Only 4 out of 31 (12.9%) hsPRS-v1 pairs detected by the union of $Y2H_{HIS3}$ and lcn$Y2H_{GFP}$ were detected with both methods, indicating a high degree of orthogonality. In addition, $Y2H_{GFP}$ implemented in this study includes additional differences such as high-copy and C-terminal fusion constructs for human proteins. Therefore, we conservatively estimate 90% orthogonality between $Y2H_{HIS3}$ and $Y2H_{GFP}$ (*i.e.* ~90% of detected interactions are different, $O_{HIS3+GFP}$= 90% ). Thus, we estimate that the fraction of all true interactions captured by our merged interactome maps is $C_{HIS3+GFP} = (C_{HIS3} + C_{GFP}) \times O_{HIS3+GFP} \cong (0.125 + 0.095) \times 0.9 = 19.7\%$.

**yN2H Validation**

To assess the quality of our dataset, we used the yN2H assay (Choi et al., 2019) for validation of PPIs detected in both $HuSCI_{HIS3}$ and $HuSCI_{GFP}$. This assay was performed as previously described (Choi et al., 2019). Briefly, viral ORFs and human interaction partners were transferred into pDEST-N2H plasmids (pDEST-N2H-N1, -N2, -C1, and -C2) containing a *LEU2* (N1/C1 vectors) or a *TRP1* (N2/C2 vectors) auxotrophy marker, via Gateway LR cloning. Plasmid DNA was extracted from bacteria using a 96-well DNA extraction kit (Qiagen, 27193) and transformed into haploid *S. cerevisiae* Y8800 (MATa) and Y8930 (MATα) strains. The resulting transformants carrying N1/C1/Fragment1 vectors (*LEU2* cassette, in Y8930 strain) or N2/C2/Fragment2 vectors (*TRP1* cassette, in Y8800 strain) were inoculated into a 96-well microplate (Fisher Scientific/Corning, 07200720A) containing 160 µL liquid SC-Leu or SC-Trp media per well, respectively. In parallel, established hsPRS-v2/hsRRS-v2 (Choi et al., 2019) were inoculated. In addition, two protein pairs from the hsPRS-v2, with different N2H signal intensities, were included in duplicates on every plate (NCBP1/NCBP2 and SKP1/SKP2). We also included a set of ~100 randomly selected pairs of viral human protein pairs as a random reference set (vhRRS; **Supplementary Table 2B**), for which human proteins were randomly selected from hsRRS-v2. Viral human protein pairs in vhRRS and other viral human datasets were randomly distributed in the plates and tested together with hsPRS-v2/hsRRS-v2, which were in separate plates.

After overnight incubation at 30°C of the haploid cultures, mating was performed by transferring 5µL of each haploid strain into a 96-well microplate containing 160 µL YEPD medium per well. The combined cells were incubated overnight at 30°C. Yeast cells expressing viral or human ORFs were also mated with a yeast strain harbouring a plasmid containing only Fragment 1 or Fragment 2 in order to measure background signal (e.g., N1-X was mated with N2-Y, where X and Y are the proteins of a tested pair, as well as Fragment 2 and vice versa). Selection for diploid yeast cells was conducted by inoculating 10 µL of mated yeast into a 96-well microplate containing 160 µL medium per well (SC-Leu-Trp). After overnight incubation at 30°C, 50 µL was transferred from the first diploid selection culture into 1.2 mL medium (SC-Leu-Trp) in deep well plates (Qiagen, 19579), which were incubated overnight at 30°C with shaking at 900 rpm. These plates were then centrifuged (2,500 rpm, 15 min) and the supernatant discarded. Each yeast cell pellet was individually resuspended in 100 µL of the NanoLuc Assay solution by gently pipetting up-and-down several times until fully resuspended. Homogenized solutions were transferred into white flat-bottom 96-well plates (Greiner Bio-One, 655073) that were then incubated in the dark (1h, room temperature). Using a luminometer (TriStar² S LB 942, Berthold, 61457) set up for one-second orbital shaking before each measurement, luminescence was evaluated for each sample (integration time of the luminescence signal was set at 2s per sample). To score each protein pair X–Y, we calculated a normalized luminescence ratio (NLR) corresponding to the raw luminescence value of the tested pair (X–Y) divided by the maximum luminescence value from one of the two controls (X-Fragment 2 or Fragment 1-Y) (Choi et al., 2019). The 1% RRS threshold was based on the vhRRS and determined using the R quantile function.

**Bioinformatic analysis of binary interactome map**

Network construction and visualization

Starting from a list of protein interaction pairs detected by $Y2H_{HIS3}$ or $Y2H_{GFP}$ and validated by yN2H, we constructed a graph (network) where viral proteins and human targets are represented by vertices (nodes), and interactions are represented by edges (links). Cytoscape (v3.8.2) was used to visualize the interaction network (Shannon et al., 2003).

Gene Ontology enrichment analysis

The Metascape web-platform was used to select GO terms enriched for our binary interaction human targets and four different AP-MS studies (Gordon et al., 2020a, 2020b; Li et al., 2020; Nabeel-Shah et al., 2020; Stukalov et al., 2020; Zhou et al., 2019). The human ORFeome collection used for the Y2H screen was used as the background for our binary interaction host proteins, and the background for four different AP-MS studies was according to their screening search space. GO terms with a hypergeometric $p < 0.01$, an observed gene count ≥ 3 and an enrichment factor >1.5 were collected and grouped into clusters based on their membership similarities. The adjusted *p*-values were calculated using the Benjamin-Hochberg procedure. The enrichment factor is the ratio between the observed and the expected counts. Kappa scores were used as the similarity metric when performing hierarchical clustering on the enriched terms, and sub-trees with a similarity > 0.3 were considered a cluster. The most statistically significant GO term within a cluster was chosen to represent the cluster.

**Identification of interologs via pairwise Y2H testing**

Curation of previously known binary interactions involving SARS-CoV-1 and host proteins

To identify interologs representative of conserved interactions, we examined known binary interactions between SARS-CoV-1 and human proteins (Orchard et al., 2014) by applying a *HIS3*-based Y2H assay individually to each pair. A curated list of these virus-host interactions (97 unique interactions) can be found in **Supplementary Table 5**. The barcoded collections described above (bhORFeome, and barcoded SARS-CoV-2 destination ORF collections) were used for these interaction tests, and the pairwise tests were performed in the same orientation as the large scale $Y2H_{GFP}$ screen (bhORFeome in pAR068 and viral ORFs in pHiDEST-DB). A total of 77 binary interactions involving barcoded human ORFs available in our collection were examined.

*Pairwise testing of known SARS-CoV-1 binary interactions to identify interologs*

Individual *E. coli* clones of barcoded human ORFs were cherry picked from our bhORFeome (in pAR068) using a BioMatrix Robot (S&P Robotics Inc.). Approximately 20% of these clones were then verified via Sanger sequencing (>90% success rate for both ORF and BC sequences). A total of 63 human destination plasmid clones (60 with 2 different unique barcode sets and 3 with a single unique barcode set) were identified for this assay. The *E. coli* clones bearing each barcode set were cultured in 96-deepwell plates with LB + 100 µg/mL carbenicillin media at 37°C for 48h. Cultures for the two barcode sets for the same ORF were merged in equal amounts before *E. coli* plasmid mini-prep (Macherey-Nagel, 740727.250). Plasmid DNA was then transformed into the RY3011 yeast strain in 96-well format to keep each ORF separate (Frozen-EZ yeast transformation II kit, Zymo Research, T2001). The resultant haploid prey (AD-Y) transformants were cultured to saturation at 30°C in SC media lacking tryptophan (SC-Trp). Glycerol stocks were made with a final concentration of 25% glycerol and then stored at -80°C. The haploid bait strains of individual virus ORFs were previously prepared (see above). Haploids were arranged in a 96-well format to accommodate the pairwise assay.

Frozen haploid glycerol stocks of both bait and prey strains were inoculated (5 uL) into 180 uL of haploid growth media (SC-Leu for bait, and SC-Trp for prey) and grown at 30°C for until saturation (48-72h). Mating was then performed individually by mixing 40 µL of saturated haploid yeast cultures in 96-well plates, followed by a 24h incubation in 2x YPAD media at room temperature. The mated cultures were then washed with dd$H_2O$, transferred to diploid selection media (SC-Leu-Trp+10xHis), and cultured at 30°C for 3 days. Next, 7 µL of saturated diploid cultures were spotted onto solid diploid selection media and incubated at 30°C for 4 days. Diploid yeast spots were then inoculated into fresh liquid diploid media and grown to saturation in deep-well plates. As a negative control, a pool of empty barcoded AD strains were also mated with the individual DB-X strains, to detect any auto-activation among the viral baits. Resultant diploids along with glycerol stocks of assay diploids were stored at -80°C.

To perform the *HIS3*-based Y2H growth-selection assay, cultures were started by inoculating 20µL of glycerol stock into a total volume of 1,200 µL liquid diploid selection media (SC-Leu-Trp+10xHis). Following growth saturation at 30°C, 100 µL of these cultures were transferred into 96-well plates. The BioMatrix robot was then used to condense 96-well plates into 384-well plates by pinning individual wells onto plates bearing solid diploid media for outgrowth at 30°C for 3 overnights. We then used the BioMatrix robot to transfer yeast colonies (now in 384-well format) to plates containing dd$H_2O$, which were subsequently used to spot yeast onto the following Y2H selection media plates in duplicates: i) diploid media, SC-Leu-Trp+10xHis, ii) -His selection, SC-Leu-Trp-His, and iii) -His+3-AT selection, SC-Leu-Trp-His+1mM 3-AT. Plates were incubated at 30°C for 3 overnights before image capture and analysis. Successful interactions were indicated by the presence or absence of colony growth in both replicates, upon selection in -His and/or in the more stringent -His+3-AT conditions.

#### QUANTIFICATION AND STATISTICAL ANALYSIS

Data in **Fig. 1C** are presented as a fraction (%) of protein pairs in a given set identified as positive at 1% positive vhRRS threshold. Error bars represent standard error of the positive fractions, *p* value was derived from Fisher's exact test. Data in **Fig. 2B** are presented as percentages of observed items over background items, *p* value was derived from hypergeometric distribution. Data in **Fig. 2C** are presented as fractions (%) of random interactors or protein interactions that were observed either in previously known human host targets or identified intraviral protein interactions. *p* value was derived from empirical testing.

#### REFERENCES

Altmann, M., Altmann, S., Falter, C., and Falter-Braun, P. (2018). High-quality yeast-2-hybrid interaction network mapping. Curr Protoc Plant Biol *3*, e20067.

Altmann, M., Altmann, S., Rodriguez, P.A., Weller, B., Elorduy Vergara, L., Palme, J., Marín-de la Rosa, N., Sauer, M., Wenig, M., Villaécija-Aguilar, J.A., et al. (2020). Extensive signal integration by the phytohormone protein network. Nature *583*, 271–276.

Braun, P., Tasan, M., Dreze, M., Barrios-Rodiles, M., Lemmens, I., Yu, H., Sahalie, J.M., Murray, R.R., Roncari, L., de Smet, A.-S., et al. (2009). An experimentally derived confidence score for binary protein-protein interactions. Nat. Methods *6*, 91–97.

Choi, S.G., Olivet, J., Cassonnet, P., Vidalain, P.-O., Luck, K., Lambourne, L., Spirohn, K., Lemmens, I., Dos Santos, M., Demeret, C., et al. (2019). Maximizing binary interactome mapping with a minimal number of assays. Nature Communications *10*, 3907.

Gibson, D.G., Young, L., Chuang, R.-Y., Venter, J.C., Hutchison, C.A., 3rd, and Smith, H.O. (2009). Enzymatic assembly of DNA molecules up to several hundred kilobases. Nat. Methods *6*, 343–345.

Gordon, D.E., Jang, G.M., Bouhaddou, M., Xu, J., Obernier, K., White, K.M., O’Meara, M.J., Rezelj, V.V., Guo, J.Z., Swaney, D.L., et al. (2020a). A SARS-CoV-2 protein interaction map reveals targets for drug repurposing. Nature *583*, 459–468.

Gordon, D.E., Hiatt, J., Bouhaddou, M., Rezelj, V.V., Ulferts, S., Braberg, H., Jureka, A.S., Obernier, K., Guo, J.Z., Batra, J., et al. (2020b). Comparative host-coronavirus protein interaction networks reveal pan-viral disease mechanisms. Science *370*, eabe9403.

Kim, D.-K., Knapp, J.J., Kuang, D., Chawla, A., Cassonnet, P., Lee, H., Sheykhkarimli, D., Samavarchi-Tehrani, P., Abdouni, H., Rayhan, A., et al. (2020). A comprehensive, flexible collection of SARS-CoV-2 coding regions. G3 *10*, 3399–3402.

Kim, D.-K., Sheykhkarimli, D., Kishore, N., Rayhan, A., Kuang, D., Li, R., Li, S., Škalič, M., Colobella, C., Cote, A.G., et al. (2021). Global dynamics of environment-dependent interactomes. Unpublished.

Kingma, D.P., and Ba, J. (2014). Adam: A method for stochastic optimization. arXiv 1412.6980.

Langmead, B., and Salzberg, S.L. (2012). Fast gapped-read alignment with Bowtie 2. Nat. Methods *9*, 357–359.

Li, J., Guo, M., Tian, X., Wang, X., Yang, X., Wu, P., Liu, C., Xiao, Z., Qu, Y., Yin, Y., et al. (2020). Virus-host interactome and proteomic survey reveal potential virulence factors influencing SARS-CoV-2 pathogenesis. Med (N Y) *2*, 99–112.e7.

Luck, K., Kim, D.-K., Lambourne, L., Spirohn, K., Begg, B.E., Bian, W., Brignall, R., Cafarelli, T., Campos-Laborie, F.J., Charloteaux, B., et al. (2020). A reference map of the human binary protein interactome. Nature *580*, 402–408.

Maas, A.L., Hannun, A.Y., and Ng, A.Y. (2013). Rectifier nonlinearities improve neural network acoustic models. In Proc. Icml, (Citeseer), p. 3.

Nabeel-Shah, S., Lee, H., Ahmed, N., Marcon, E., Farhangmehr, S., Pu, S., Burke, G.L., Ashraf, K., Wei, H., Zhong, G., et al. (2020). SARS-CoV-2 Nucleocapsid protein attenuates stress granule formation and alters gene expression via direct interaction with host mRNAs. bioRxiv 342113.

Orchard, S., Ammari, M., Aranda, B., Breuza, L., Briganti, L., Broackes-Carter, F., Campbell, N.H., Chavali, G., Chen, C., del-Toro, N., et al. (2014). The MIntAct project--IntAct as a common curation platform for 11 molecular interaction databases. Nucleic Acids Res. *42*, D358–D363.

Shannon, P., Markiel, A., Ozier, O., Baliga, N.S., Wang, J.T., Ramage, D., Amin, N., Schwikowski, B., and Ideker, T. (2003). Cytoscape: a software environment for integrated models of biomolecular interaction networks. Genome Res. *13*, 2498–2504.

Stukalov, A., Girault, V., Grass, V., Bergant, V., Karayel, O., Urban, C., Haas, D.A., Huang, Y., Oubraham, L., Wang, A., et al. (2020). Multi-level proteomics reveals host-perturbation strategies of SARS-CoV-2 and SARS-CoV. bioRxiv 156455.

The ORFeome Collaboration (2016). The ORFeome collaboration: a genome-scale human ORF-clone resource. Nature Methods *13*, 191–192.

Weile, J., Sun, S., Cote, A.G., Knapp, J., Verby, M., Mellor, J.C., Wu, Y., Pons, C., Wong, C., van Lieshout, N., et al. (2017). A framework for exhaustively mapping functional missense variants. Mol. Syst. Biol. *13*, 957.

Wu, A., Peng, Y., Huang, B., Ding, X., Wang, X., Niu, P., Meng, J., Zhu, Z., Zhang, Z., Wang, J., et al. (2020a). Genome composition and divergence of the novel coronavirus (2019-nCoV) originating in China. Cell Host & Microbe *27*, 325–328.

Wu, F., Zhao, S., Yu, B., Chen, Y.-M., Wang, W., Song, Z.-G., Hu, Y., Tao, Z.-W., Tian, J.-H., Pei, Y.-Y., et al. (2020b). A new coronavirus associated with human respiratory disease in China. Nature *579*, 265–269.

Yachie, N., Petsalaki, E., Mellor, J.C., Weile, J., Jacob, Y., Verby, M., Ozturk, S.B., Li, S., Cote, A.G., Mosca, R., et al. (2016). Pooled‐matrix protein interaction screens using Barcode Fusion Genetics. Mol. Syst. Biol. *12*, 863.

Yu, F., and Koltun, V. (2015). Multi-scale context aggregation by dilated convolutions. arXiv 1511.07122.

Yu, H., Braun, P., Yildirim, M.A., Lemmens, I., Venkatesan, K., Sahalie, J., Hirozane-Kishikawa, T., Gebreab, F., Li, N., Simonis, N., et al. (2008). High-quality binary protein interaction map of the yeast interactome network. Science *322*, 104–110.

Zhou, Y., Zhou, B., Pache, L., Chang, M., Khodabakhshi, A.H., Tanaseichuk, O., Benner, C., and Chanda, S.K. (2019). Metascape provides a biologist-oriented resource for the analysis of systems-level datasets. Nat. Commun. *10*, 1523.
